# Legume genome structures and histories inferred from *Cercis canadensis* and *Chamaecrista fasciculata* genomes

**DOI:** 10.1101/2024.09.03.611065

**Authors:** Hyun-oh Lee, Jacob S Stai, Qiaoji Xu, Thulani Hewavithana, Rabnoor Batra, Alex Liu, Brandon D Jordan, Rachel Walstead, Jerry Jenkins, Melissa Williams, Jenell Webber, Jane Grimwood, John T Lovell, Tomáš Brůna, Shengqiang Shu, Keykhosrow Keymanesh, Joanne Eichenberger, Jeremy Schmutz, David M Goodstein, Kerrie Barry, David Sankoff, Lingling Jin, James H Leebens-Mack, Steven B Cannon

**Affiliations:** ORISE Fellow, USDA-ARS, Corn Insects and Crop Genetics Research Unit, 819 Wallace Rd, Ames, IA 50011, USA; Department of Mathematics and Statistics, University of Ottawa, Ottawa, Ontario K1N 6N5, Canada; Department of Computer Science, University of Saskatchewan, Saskatoon, Saskatchewan S7N 5C9, Canada; School of Computer Science, University of Waterloo, Waterloo, Ontario N2L 3G1, Canada; USDA-ARS, Corn Insects and Crop Genetics Research Unit, 819 Wallace Rd, Ames, Iowa, USA; Genome Sequencing Center, HudsonAlpha Institute for Biotechnology, Huntsville, Alabama 35806, USA; US Department of Energy Joint Genome Institute, Berkeley, California 94720, USA; Department of Biology, University of Georgia, Athens, Georgia 30602, USA

**Keywords:** allopolyploidy, Caesalpinoideae, Cercidoideae, *Cercis canadensis* (redbud), *Chamaecrista fasciculata* (partridge pea), Fabaceae, legumes, whole genome duplication

## Abstract

- The legume family originated ca. 70 million years ago and soon diversified into at least six lineages (now extant subfamilies). The signal of whole genome duplications (WGD) is apparent in species sampled from all six subfamilies. The early diversification has posed difficulties for resolving the legume backbone structure and the timing of WGDs.
- In this study, we report the genome sequences and annotations for *Cercis canadensis* (Cercidoideae) and *Chamaecrista fasciculata* (Caesalpinoideae) to help resolve the relative taxonomic placements along the legume backbone, the timings of WGDs relative to subfamily origins, and the ancestral legume karyotype.
- Analyses of genome assemblies from four subfamilies within Fabaceae show that the last common ancestor of all legumes likely had seven chromosomes, with a genome structure similar to the extant *Cercis* genome. Our analysis supports an allopolyploid origin of the subfamily Caesalpinoideae, with progenitors involving lineages along the backbone of the legume phylogeny.
- A probable allopolyploid origin of Caesalpinoideae subfamily provides a partial explanation for the difficulty in resolving the structure of the legume backbone. The retained karyotype structure and lack of a WGD in the last 100+ Mya, underscore the utility of the *Cercis* genome as an ancestral reference for the legume family.

## Introduction

The legume family (Leguminosae or Fabaceae) is the third largest family of flowering plants, comprising about 770 genera and 19,500 species (Lewis *et al*., 2013; Azani *et al*., 2017). Species such as peanuts, cowpeas, soybeans, and fava beans are an important source of protein for a large proportion of the global human population, and play an important agricultural role in fixing nitrogen (N_2_) from the atmosphere and converting it to reduced forms that are used by plants to produce amino acids and other biomolecules (Martins *et al*., 2003; Chu *et al*., 2004; Herridge *et al*., 2008; Salvagiotti *et al*., 2009; Köpke & Nemecek, 2010).

The legume family originated within the Rosid II angiosperm clade toward the end of the Mesozoic era, around 70 Million Years Ago (MYA) (Li *et al*., 2019). Within a span of roughly 15 MY from its origin, the family had radiated giving rise to six lineages that are recognized as subfamilies: Cercidoideae, Detarioideae, Dialioideae, Duparquetioideae, Caesalpinioideae, and Papilionoideae (Azani *et al*., 2017). Most agronomically important species (and approximately two thirds of species in the family) fall within the Papilionoid clade, but the other subfamilies contain many species of great importance in term of ecological service and economic value, including timber species (e.g. *Gleditsia*, *Robinia*), forage (e.g. *Acacia*, *Caesalpinia*), and human consumption (e.g. *Tamarindus* [tamarind] *Detarium* [sweet detar], *Ceratonia* [carob], *Tylosema* [marama bean]) (Cannon *et al*., 2011).

Rapid bursts of diversification within the family have made resolution of the legume phylogeny difficult. Further complicating the understanding of the genomic evolution in the family is the presence of multiple, apparently independent whole genome duplications within most subfamilies (Cannon *et al*., 2015; Zhao *et al*., 2021). Chromosome numbers in the family vary widely, though there are clear modal counts for each subfamily: n=12 for Detarioideae, 14 for Cercidoideae, 14 for Dialoideae, and 14 for Caesalpinioideae. In Papilionoideae, the range of chromosome numbers is broad, but the majority of species in this subfamily have counts of n=7-11 (Ren *et al*., 2019).

Here, we describe high-quality genome assemblies and annotations for *Cercis canadensis* L. and *Chamaecrista fasciculata* (Michx.) Greene. These species represent two of the six generally-recognized legume subfamilies, and thus are well placed for helping to infer early events in the evolution of the legumes. We test the extent to which the *C. canadensis* genome, representing an exceptional legume lineage that has not experienced a WGD over its evolutionary history subsequent to the ~135 Mya gamma triplication (Jiao *et al*., 2012), retains pre-legume ancestral genome structure including the ancestral monoploid karyotype count for all members of the family.

For analyses of chromosome structural and gene family evolution, we make comparisons among six sequenced legume genomes that represent the four largest legume subfamilies; and also against four nonlegume outgroup species. Those outgroup species, in order from youngest to oldest shared ancestry with the legumes, are *Quillaja saponaria* (Quillajaceae, the closest family to the legumes), *Prunus persica* (Rosaceae, in the Fabideae with the legumes), *Arabidopsis thaliana* (Malvideae, sister to the Fabideae and within the Rosid clade), and *Vitis vinifera*, (Vitaceae, Vitales, sister to clade with all other orders in the Rosid clade).

Of the newly sequenced genomes reported here, *C. canadensis*, also known as the eastern redbud, is a deciduous ornamental tree native to eastern North America. The *Cercis* genus is eponymous for the Cercidoideae subfamily. There are approximately 10 species within the genus - three native to North America, six native to China and south-central Asia, and one native to the Mediterranean region (Davis *et al*., 2002). We compare the assembly of *C. canadensis* to a genome assembly of *C. chinensis* (Li *et al*., 2023), and evaluate the *C. canadensis* assembly and annotations in the context of other legume species.

*Chamaecrista fasciculata*, commonly known as partridge pea, is in the Caesalpinioideae legume subfamily. *C. fasciculata* is an annual plant common in prairies of eastern and central North America (Fenster, 1991; Govaerts *et al*., 2021). *C. fasciculata* exhibits symbiotic nitrogen fixation (SNF), hosting nitrogen-fixing Bradyrhizobium bacteria in specialized structures (nodules) on roots. *C. fasciculata* has been used as a model for research on the ecology, physiology, and evolution of symbiotic nitrogen-fixation (Singer *et al*., 2009). SNF is found in most genera in the Papilionoideae and in several clades within the Caesalpinioideae, but in none of the other legume subfamilies. SNF presence and absence among legume subfamilies has informed inference of SNF evolution with the Rosid Nitrogen Fixing Clade (Sprent *et al*., 2017; Griesmann *et al*., 2018; Zhao *et al*., 2021).

## Materials and Methods

### Cercis canadensis and Chamaecrista fasciculata genome assembly and annotation

Genome assembly and annotation methods for *Cercis canadensis* and *Chamaecrista fasciculata* are described in Supporting Information Methods S1.

### Phylogenomic analyses

For phylogenomic analyses (Figs 4, 5), legume-focused gene families were first constructed using the Pandagma gene family workflow (https://github.com/legumeinfo/pandagma; Cannon *et al*., 2024), using the CDS and protein sequences of 36 legume species in 21 genera (Table S1) and four nonlegume outgroup genera. Inputs for the initial base families included 15 individual legume species (Table S1) and exemplar sequences for pangene sets for six legume genera for which multiple annotations and species are available (*Arachis, Cicer, Glycine, Medicago, Phaseolus*, and *Vigna*). For example, the *Vigna* pangene set was calculated based on seven accessions of *V. unguiculata*, two of *V. radiata*, and one of *V. angularis*. The non-legume outgroup species were *Quillaja saponaria, Prunus persica, Arabidopsis thaliana*, and *Vitis vinifera*. There were 39,981 gene families generated using these inputs, and 25,271 families containing at least 4 sequences and at least 2 distinct genera. These base gene families are available at https://data.legumeinfo.org/LEGUMES/Fabaceae/genefamilies/legume.fam3.VLMQ/.

Using on those initial “base” families, the 15 annotation sets described in this manuscript (11 legume genera and 4 outgroup genera) were placed into the base gene families above by homology, using the “pandagma fsup” workflow, parameterized with identity >= 30% and coverage >= 40%. Protein sequences from each family were aligned using famsa (Deorowicz *et al*., 2016). The alignments were modeled using hmmbuild from the hmmer package (Finn *et al*., 2011) to generate hidden Markov models (HMMs). The original gene family sequences were then realigned to the HMM for each family, and then trimmed to the match-states of the HMM. Phylogenetic trees were then calculated using FastTree (v. 2.1)(Price *et al*., 2010). Selected gene trees (e.g. Figs. 4b-e) were also calculated using RAxML, with 1000 bootstraps.

To calculate a consensus gene phylogeny (Fig. 4a), 50 gene families were selected at random from “complete” families - i.e., families containing the expected number of genes from each species under the assumption of retention following known WGDs and no additional WGDs. Homoeologous genes from ancestrally tetraploid species in each of the 50 selected phylogenies were labeled A or B based on their position in the gene trees; for example, Medicago.A and Phaseolus.A were identified in one clade and Medicago.B and Phaseolus.B in another clade. The A and B labeling was applied top to bottom in rooted trees that had been ordered relative to the outgroup, by the Order function in the Archaeopteryx tree viewer (Han & Zmasek, 2009). This places sister clades with more nodes above those with fewer nodes, which generally results in the better-represented Papilionoid clade placed at the top, and Cercidoid clade near the bottom. The effect should otherwise be neutral with respect to other aspects of clade arrangements. Given these labeled gene families, a supermatrix alignment was generated by concatenating alignments from all 50 gene families, with genes and paralogs placed in consistent order: Lotus.A first, Medicago.A second, etc. The alignment was sampled at modulo 5 (taking every fifth amino acid) to make phylogenetic calculations tractable. The resulting alignment matrix had 9483 sites. The consensus phylogeny was then calculated using RAxML-NG (Kozlov *et al*., 2019), using the raxml-ng --all, with model LG+G8+F.

### Synonymous-site (Ks) calculations and divergence time estimation

The Ks values (Fig. 5) were calculated as part of the pandagma gene family workflow (Cannon *et al*., 2024). For a given species pair, gene pairs were identified first based on homology using mmseqs2 (Steinegger & Söding, 2017), then filtered based on inclusion in synteny blocks identified by DAGChainer (Haas *et al*., 2004). Given those gene pairs, Ks values were calculated using PAML (Yang, 2007). A modal value was then calculated for all genes in a synteny block. The modal values for the genes in the block were applied to all genes in that block, which were in turn used to calculate genome-wide Ks-value frequency distributions (Fig. 5). Modal Ks values for each species pair (both speciation and WGD peaks) were used as parameters in a system of equations to solve for each branch length in the consensus gene tree described above (Fig. 5d). Divergence times for the tree in Fig. 6a were approximated on the consensus tree using calibrations from TimeTree database (http://TimeTree.org)(Kumar *et al*., 2017). The median value used for *Vitis* and *Medicago* was taken as 117 Mya (Wikström *et al*., 2001).

### Analysis of gene family expansions and contractions and gene ontology enrichment

Homologous gene clustering was performed with the OrthoFinder (Emms & Kelly, 2019) clustering algorithm and default options (e-value 1e-2, inflation value 1.5) by the Orthovenn3 web server (Sun *et al*., 2023). Gene family contraction and expansion analysis was performed using CAFE5 (Mendes *et al*., 2020). Gene ontology (GO) terms for biological process, molecular function, and cellular component categories and enrichment of the expanded and contracted genes were then obtained. The enriched horizontal bar plot was drawn by using SRplot (Tang *et al*., 2023).

### Analysis of duplicated subgenomes resulting from polyploid events

*Chamaecriasta, Senna,* and *Bauhinia* all exhibit duplicated syntenic regions compared to *Cercis* due to their independent WGD events after diverging from *Cercis*. The SyntenyLink algorithm (Hewavithana *et al*., 2023) was used to identify two subgenomes derived from ancient allotetraploidy events in the ancestry of *Chamaecriasta, Senna,* and *Bauhinia*. SyntenyLink assesses differences in substitution and fractionation patterns in synteny blocks as homoeologous gene duplicates (syntelogs) diverge. The results are visualized using SynVisio (Bandi & Gutwin, 2020) showing syntenic blocks linking regions of the *Cercis* genome to homoeologous regions of the in *Bauhinia* (Fig. 1b), *Chamaecriasta* (Fig. 1c), and *Senna* (Fig. 1d) genomes.

**Fig. 1.**
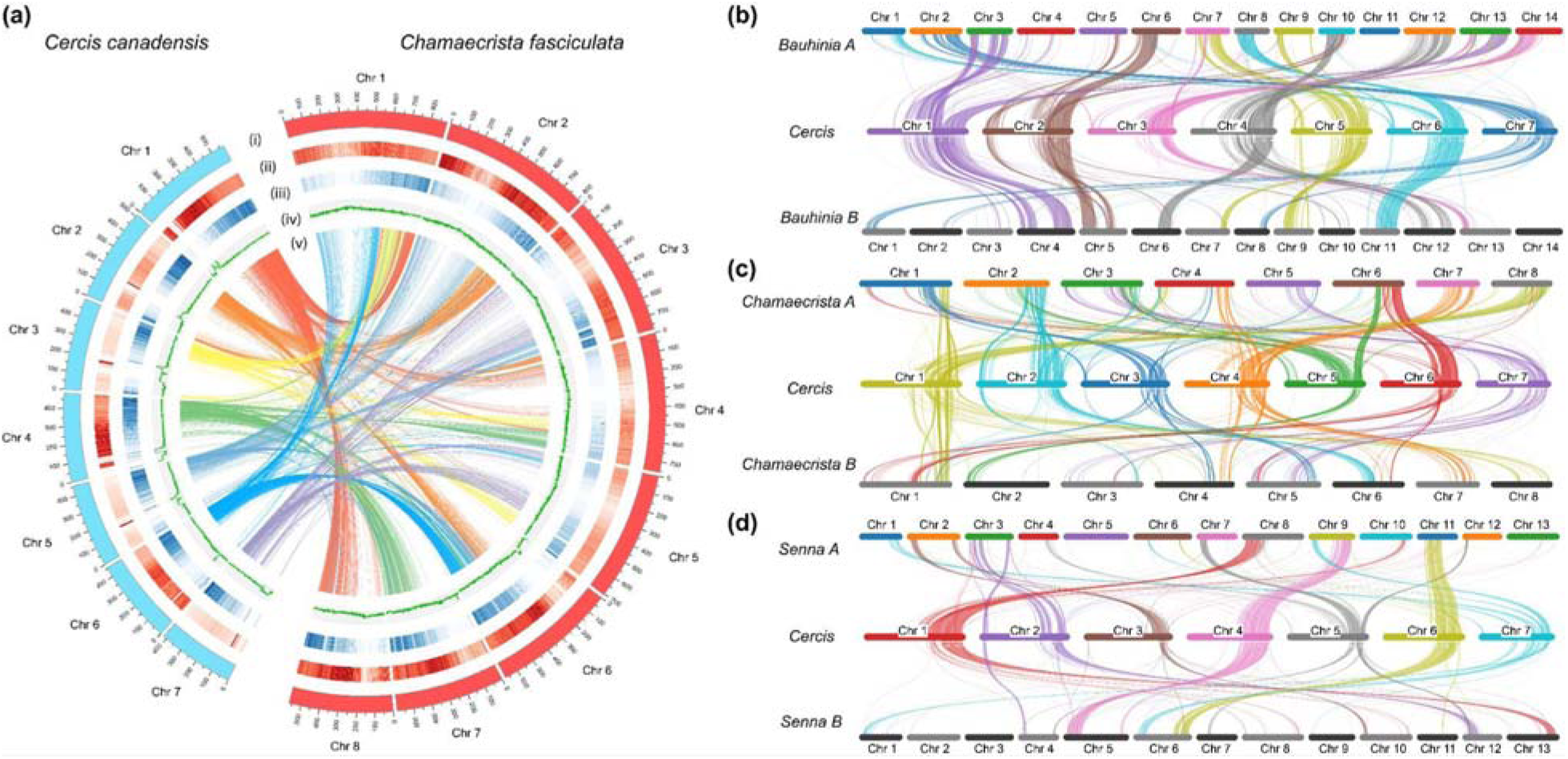
Plots of genomic features and syntenic relationships. (a) Circos plots representing features of the *C. canadensis* and *C. fasciculata* genomes. Circos track (i): chromosome length (Mbp); track (ii): repeat density, track (iii): gene density, track (iv): GC plot, inner connecting lines (v): synteny regions, identified with MCScanX. (b-d) Synteny plots shown in SyntenyLink (Bandi & Gutwin, 2020; Hewavithana *et al*., 2023) images, showing two subgenomes in *Bauhinia* (b), *Chamaecrista* (c), and *Senna* (d) relative to *Cercis* at the center of each panel and the least/most fractionated subgenome above/beneath *Cercis*.

### Ancestral genome reconstructions and rearrangement distances support seven protochromosomes

We used the RACCROCHE (Xu *et al*., 2020) procedure to infer gene content and order for hypothesized ancestral chromosomes in the genomes in a phylogeny. First, we identify adjacent genes (Yang & Sankoff, 2011), allowing up to a specified number of spacer genes between genes scored as adjacent, across the chromosomes in the input genomes. Considering the divergence times among our focal genomes, we allow 2 spacer genes to score gene adjacencies. Then, for each ancestral node in the species phylogeny, we infer adjacencies by generating graphs with all phylogenetically informative adjacencies. In the graph, vertices are adjacencies, and edges join any two adjacencies that each contain one of the 5’ and 3’ ends of the same gene. The graph is analyzed using the Maximum Weight Matching algorithm, which produces linear ancestral “contigs” as output, with each contig containing a collection of genes found in proximity in more than one of the input genomes. To avoid biases due to widely disparate contig lengths, we cut each contig into “20-mers” of length at most 20 genes, creating a larger set of more comparable contigs.

To group the reduced contigs (20-mers) into collections representing inferred ancestral chromosomes, we match them against the chromosomes of the input genomes and count the number of times any two contigs match the same chromosome. Contig ordering is taken into account, and multiple matches within a genome are permitted (to permit modeling of WGDs). This scoring produces a co-occurrence matrix, which is then clustered using a complete-linkage clustering of the contigs. The output can be interpreted as the inferred gene content of each ancestral chromosome. The reconstructed ancestors each contain at most one member of each gene family from ortholog groups calculated from the descendant genomes. We then performed “g mer” analysis (Xu *et al*., 2023) to estimate the ancestral number of chromosomes, x.

### DCJ distance inference of species relationships

The Double Cut and Join (DCJ) distance (Yancopoulos *et al*., 2005) is used to quantify the structural differences between two genomes. Smaller DCJ values indicate fewer rearrangements between the two genomes. We calculated the total number of DCJ distances between all pairs of ancestors using the UniMOG tool (Hilker *et al*., 2012).

## Results

### Genome assembly and annotation assessment

The genomes of both *C. canadensis* and *C. fasciculata* were initially assembled based on PacBio ccs data, and then the contigs were oriented and ordered using Hi-C data. Heterozygous snp/indel phasing errors were corrected using PacBio and Illumina data. The 99.46-99.58% of the assembled sequences were assigned to chromosomes. Finally, *C. canadensis* was assembled with 7 pseudochromosomes and *C. fasciculata* with 8 pseudochromosomes. Two haplotype assemblies and annotations were resolved for each species. The final chromosome-level assemblies for*C. canadensis* and *C. fasciculata*, exhibited length variation between haplotypes (Table 1). To assess the completeness of the genome, we performed BUSCO (v. 5.4.5) analysis in gene mode using the fabales_odb10 dataset and found completeness percentages of 96.4 and 93.6% (single copy percentages of 92.3 and 79.1%). The missing rates were 3.4% and 6.1% (Table 1). The relatively high “missing” BUSCO rates are likely due to the fact that fabales_odb10 consisted of only 10 species in the Papilionidae family, and when we expanded the BUSCO DB to eudicots_odb10, *C. canadensis* and *C. fasciculata* were found to have 99.4% and 99.5% complete BUSCOs, respectively.

**Table 1.** Genome and gene statistics for *Cercis canadensis* and *Chamaecrista fasciculata*.

| Species | *Cercis canadensis* |  | *Chamaecrista fasciculata* |  |
| --- | --- | --- | --- | --- |
| Haplotype | 1 | 2 | 1 | 2 |
| Pseudochromosome number | 7 | 7 | 8 | 8 |
| Total scaffold length (bp) | 342,014,377 | 315,499,427 | 580,459,023 | 564,372,522 |
| No. of scaffolds | 43 | 7 | 24 | 17 |
| No. of contigs | 56 | 20 | 110 | 107 |
| N50 of scaffolds (bp) | 48,286,772 | 43,568,431 | 75,709,764 | 71,910,870 |
| N50 of contigs (bp) | 27,822,107 | 26,270,282 | 11,464,191 | 9,553,364 |
| GC Ratio (%) | 36.31 | 35.75 | 35.35 | 35.22 |
| No. of gene | 27,440 | 26,713 | 29,074 | 28,859 |
| No. of cds | 51,848 | 51,022 | 49,343 | 49,167 |
| No. of exon | 360,595 | 359,144 | 329,134 | 329,851 |
| Avg. exons per cds | 6.4 | 6.4 | 6.2 | 6.2 |
| Avg. gene length (bp) | 4,047 | 4,094 | 4,257 | 4,281 |
| Avg. cds length (bp) | 1,546 | 1,553 | 1,465 | 1,471 |
| Avg. exon length (bp) | 324 | 322 | 315 | 314 |
| Total gene length (bp) | 111,055,993 | 109,386,083 | 123,788,266 | 123,559,672 |
| Total cds length (bp) | 80,207,802 | 79,275,015 | 72,295,815 | 72,355,461 |
| Longest cds (bp) | 16,539 | 16,539 | 16,455 | 16,461 |
| Shortest cds (bp) | 93 | 75 | 93 | 96 |
| Complete BUSCOs (C) | 5175 / 96.4% | 5106 / 95.2% | 5023 / 93.6% | 5018 / 93.5% |
| Complete and single-copy BUSCOs (S) | 4954 / 92.3% | 4943 / 92.1% | 4242 / 79.1% | 4243 / 79.1% |
| Complete and duplicated BUSCOs (D) | 221 / 4.1% | 163 / 3.0% | 781 / 14.6% | 775 / 14.4% |
| Fragmented BUSCOs (F) | 10 / 0.2% | 23 / 0.4% | 15 / 0.3% | 17 / 0.3% |
| Missing BUSCOs (M) | 181 / 3.4% | 237 / 4.4% | 328 / 6.1% | 331 / 6.2% |
| Total BUSCO groups searched | 5,366 | 5366 | 5,366 | 5,366 |

### Comparisons of the *Cercis canadensis* and *Chamaecrista fasciculata* genomes with resources from related species

The *C. canadensis* genome assembly described here is the second near-complete assembly published from this genus. Comparisons with the assembly for *C. chinensis* (Li *et al*., 2023) show the two assemblies to be similar in size and structure, but with large rearrangements on chromosomes 3 and 5 (Fig. S2), and having an average identity in alignable regions of only 93.76%. The assembly sizes for the two species are similar: 352.8 and 342.0 Mbp for the total assembly sizes in *C. chinensis* and *C. canadensis*; and 331.8 and 340.3 Mbp for the chromosomally anchored sequences in *C. chinensis* and *C. canadensis*.

The *C. fasciculata* genome assembly described here is the second assembly of this species – the first being a contig-level assembly (Griesmann *et al*., 2018), for isolate NF-2018-5 (derived from line MN87), GenBank accession GCA_003254925.1, with scaffold N50 of 56.6 kb and total assembly length of 429.1 Mb. The chromosome-scale, haplotype-resolved assembly described here, of isolate ISC494698, has scaffold N50 of 757.1 kb and total assembly length of 580.4 Mb.

### Synteny relationships show independent WGDs early in four legume subfamilies, but excluding *Cercis*

Synteny plots (Figs 1, 2) show a general 1::2 pattern of chromosomal duplication between *Cercis* and other legume and close outgroup species. This pattern can be seen, for example, in *Cercis* chromosome 1 matching *Bauhinia* chromosomes 3 and 4 (Fig. 1b), *Chamaecrista* chromosomes 1 and 8, (Figs 1a, c), and *Senna* chromosomes 8 and 13 (Fig. 1c). These patterns are consistent with WGDs occurring independently in the two subfamilies (*Cercis* and *Bauhinia* in the Cercidoideae subfamily, and *Chamaecrista* and *Senna* in the Caesalpinioideae subfamily), but after divergence of *Cercis* from the remaining species in the Cercidoideae (though see discussion below regarding inference of allopolyploidy in both subfamilies).

**Fig. 2.**
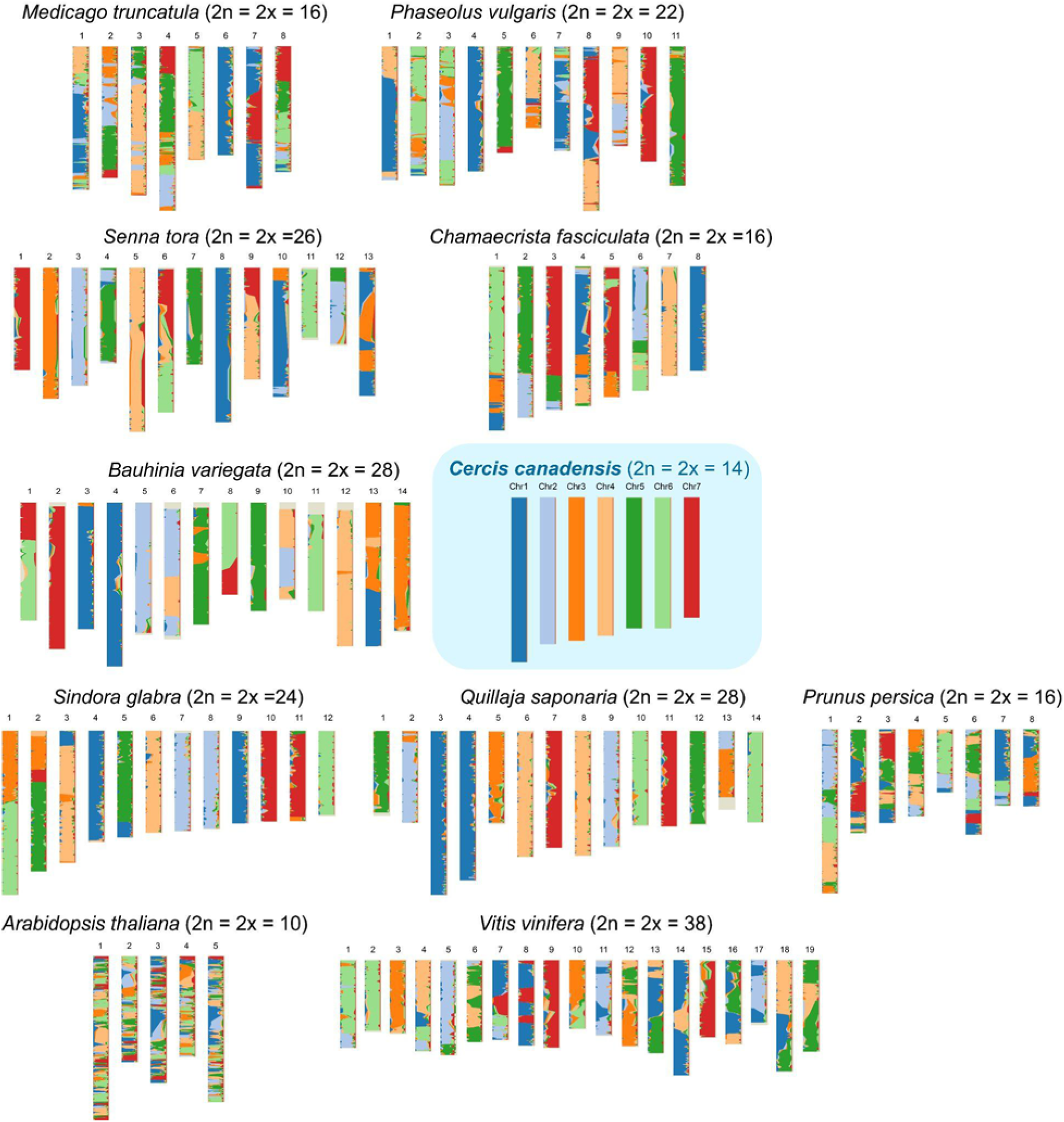
Synteny map of *Cercis canadensis* and comparison against legume and outgroup species. Chromosome-level synteny maps were generated with the PanSyn program using *Cercis* as a reference relative to each indicated comparison species.

Comparisons of *Cercis* to itself (Fig. S1) identifies only small synteny blocks and no evidence of a recent whole genome duplication. The fragmentary duplications in the *Cercis* self-comparison, together with *Ks* results (presented below) are consistent with a WGD in the timeframe of the ~135 Mya gamma triplication (Jiao *et al*., 2012).

### Inference of a seven-chromosome legume progenitor with genome structure similar to that of *Cercis*

Synteny and shared-contig analyses suggest that the *Cercis* genome structure is similar to the ancestral legume karyotype, which we infer was also likely to have had seven chromosomes. The *Quillaja, Sindora, Bauhinia,* and *Senna* chromosome structures can all be represented in terms of doublings of the seven-chromosome *Cercis* genome, with a small number of rearrangements in each lineage. In *Quillaja* (a close outgroup to the legumes), the chromosomal correspondence with *Cercis* is a simple 1::2 match, with *Cercis* chromosomes generally matching two *Quillaja* chromosomes – often with chromosomal-scale synteny across both species. The taxa requiring the fewest rearrangements relative to a simple doubling of *Cercis* (1n=7) are *Quillaja* (1n=14), requiring approximately 3 rearrangements; *Sindora* (1n=12), requiring approximately 6 rearrangements; and *Senna* (1n=13), requiring approximately 10 rearrangements. *Medicago* (1n=8) and *Phaseolus* (1n=11) show more complex restructuring - which would be consistent with the reduced chromosome counts from a hypothesized Papilionoid progenitor with 1n=14 chromosomes.

Analysis of contig co-occurrence among reconstructed ancestral genomes (Fig. 3) supports an ancestral legume karyotype with 7 chromosomes, with little rearrangement relative to current *Cercis* chromosomes. In the heat maps in Fig. 3, syntenic contigs identified across the 10 indicated species (7 legumes and 3 outgroups) are clustered by proximity, following three hypothetical phylogenetic topologies. The clusters indicate probable ancestral chromosomes, as they show syntenic contig groups that are found in proximity, at the indicated phylogenetic node, as assessed within the included species and ancestors. The heat maps correspond with the legume ancestors, supporting the estimated 7-chromosome karyotype.

**Fig. 3.**
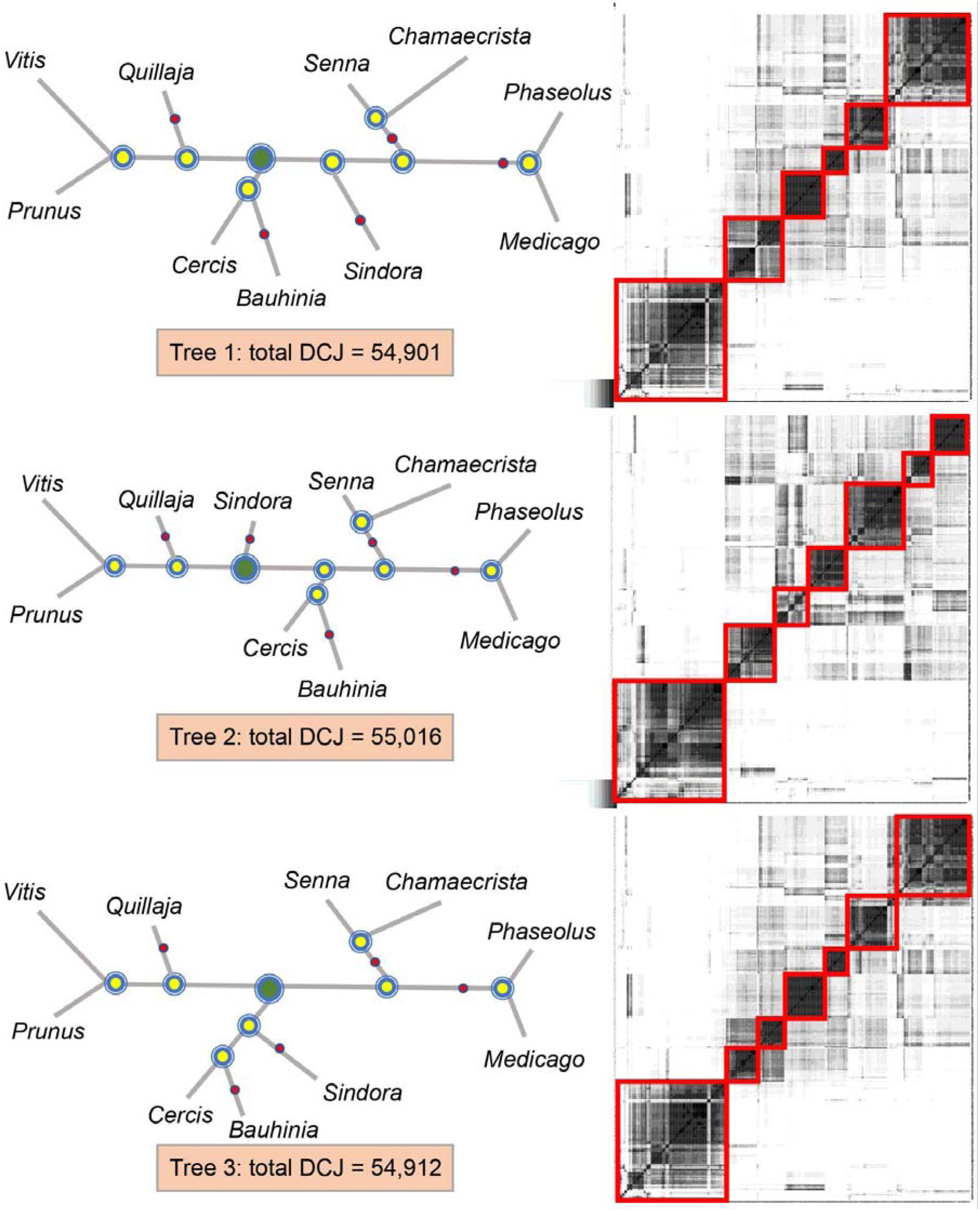
Hypothetical phylogenetic relationships of 10 selected legume and dicot outgroup species, and heat maps of contig adjacencies representing inferred ancestral chromosome content. Left: three hypothesized phylogenetic relationships. The legume ancestor node is highlighted in green while the other ancestors are in yellow. The total Double Cut and Join (DCJ) distances between ancestral nodes suggest Tree 1 as the most likely of these phylogenies. Right: Heat maps of the legume ancestors of the corresponding phylogenetic relationships (left side), showing the clusters of reconstructed contigs likely making up either six or seven ancestral chromosomes.

Fig. 3 shows seven clusters of the inferred legume ancestor formed from the co-occurrence matrix. Given the hypothesized contig content of each chromosome, the contigs are ordered on a chromosome using a Linear Ordering Problem routine. We find that the monoploid number of the legume family is likely to be x = 7 (Fig. 3), and that Tree 1 is the most parsimonious hypothetical phylogeny.

### Phylogenomic analyses and a consensus phylogenetic model of speciation and whole genome duplications

The duplication and speciation histories for the legumes in this study are evident in gene families constructed from the proteomes from each species. Using eleven representative genome sequences and annotations from four legume subfamilies as well as selected nonlegume outgroups, we calculated gene families for all genes. From 50 gene families with no sequence losses or gains relative to known WGD events, we calculated a consensus gene family based on a concatenated alignment from those families (Fig. 4a), which shows relative placements of speciation and WGD events. The basic species topology is congruent with the legume backbone topology that has been reported previously (Azani *et al*., 2017; Ferreira, 2024). Separate WGDs are apparent early in each of four legume subfamilies included in this study, but complexities in the Cercidoideae and Caesalpinioideae require discussion and interpretation.

**Fig. 4.**
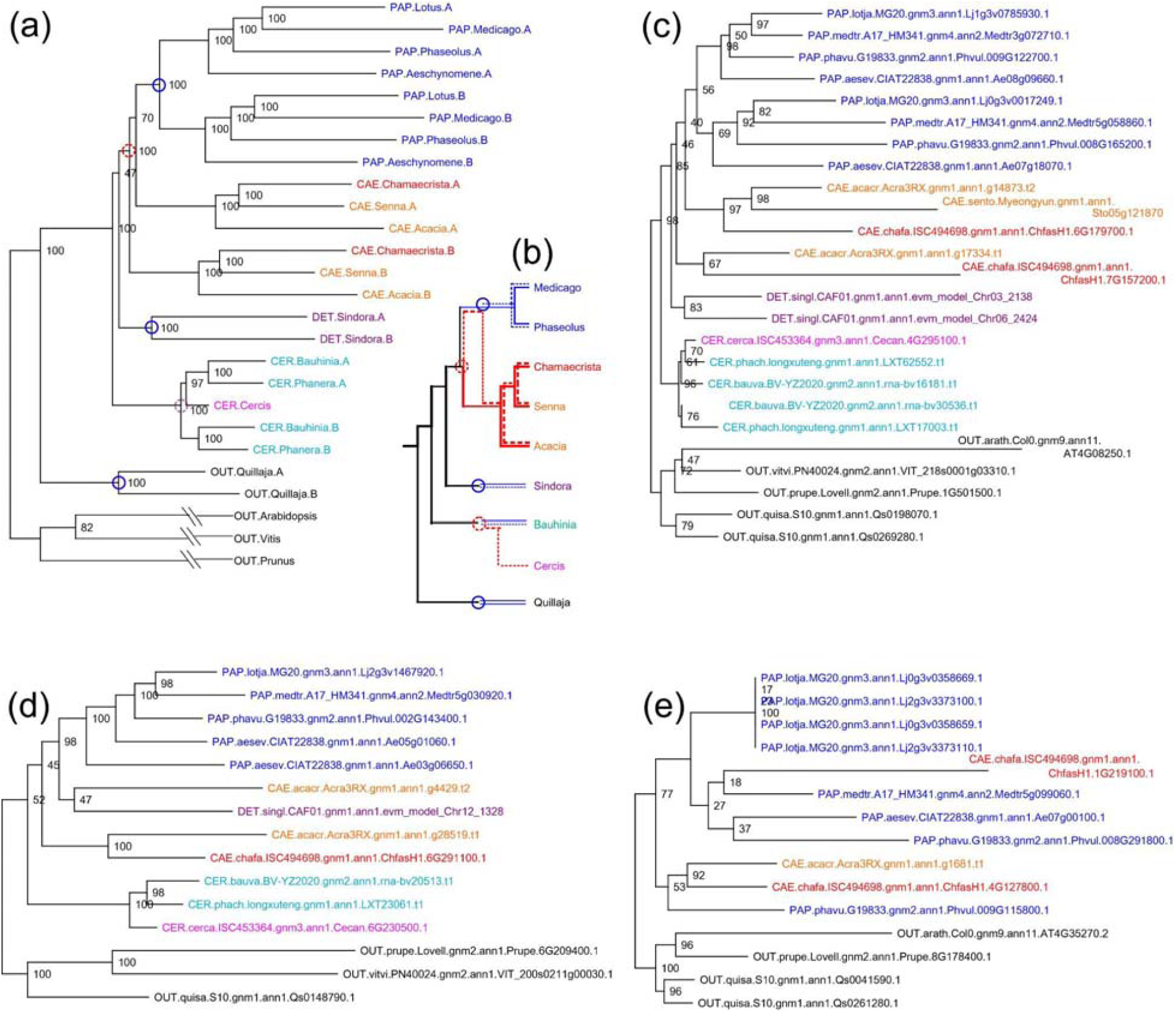
Consensus and example gene family trees. (a) Consensus tree showing inferred whole genome duplications (WGDs), calculated from 50 gene families with near-complete representation of species and WGD-derived gene duplications. Red circles indicate locations of inferred WGD events. Circles: inferred polyploidy events (blue auto-, red allo-polyploidy). (b) Schematic of gene, species, duplication paths, showing inferred allo- and auto-polyploid events. Individual lines represent evolutionary paths between genes, blue and red indicating inferred auto- and allo-polyploidy respectively. Dotted and solid lines show hypothetical alternate paths for genes from different subgenomes; for example, the path between two *Chamaecrista* paralogs would consist of a dotted path, with greater similarity to the Papilionoideae and a solid-red path with greater similarity to an older progenitor. (c) Gene family phylogeny for the family of MtNSP2 (Nodulation Signaling Pathway 2), showing the same structure as the consensus tree. (d) Gene family phylogeny of the family of LjSYMRK/MtDMI2 (SYMbiotic Receptor Kinase / Doesn’t Make Infections 2), showing loss of orthologs from non-nodulating Senna and presence in the included nodulating species. (e) Gene family phylogeny of the family of LjNIN (Nodule Inception), showing presence in all included nodulating species and absence from all non-nodulating species.

In the Cercidoideae, WGDs clearly affect *Bauhinia* and *Phanera* but not *Cercis*. However, in the consensus gene phylogeny (as well as in many individual gene phylogenies examined), *Cercis* groups with one of the *Bauhinia/Phanera* post-duplication lineages rather than outside (and prior to) the inferred WGD. A plausible explanation for this topology is that the WGD in the Cercidoideae was allopolyploid in nature, with a progenitor of *Cercis* contributing one subgenome and some other species early in the origin of the family contributing the other subgenome in the allopolyploid event.

In the Caesalpinioideae, a WGD is clearly evident, resulting in paralogs in each species examined in this subfamily; yet confusingly, one WGD-derived lineage groups more frequently with Papiloinoid species than with the Caesalpinioid paralogs. As with the Cercidoideae, allopolyploidy is a plausible explanation for this pattern. Specifically, the observed consensus gene family topology is consistent with a speciation along the legume backbone followed by a significant period of divergence (several million years), followed by an allopolyploid merger that resulted in the Caesalpinoid subfamily – each member of which would have two (divergent) subgenomes. One of the diploid lineages would then have then gone on to become the progenitor of the Papilionoideae - which experienced its own WGD prior to diversification within that subfamily.

The hypothesized allopolyploid origins of the Caesalpinioideae and the Cercidoideae can be seen in the schematic in Fig. 4b. Here, an early speciation in the lineage leading to the Caesalpinioideae and Papilionoideae is represented as a red circle. After significant divergence, merger of two descendant species would have given rise to the 1n=14 Caesalpinioideae. In any species within that subfamily, two subgenomes are present, and paralogs from those two subgenomes may have distinct evolutionary histories that reflect the different histories of the two species that contributed to the fundamentally allopolyploid subfamily. In the schematic, those distinct gene histories can be seen as distinct dotted and solid paths leading to the speciation origin (red circle).

Similar dual paths can be seen in the Fig. 4b schematic of allopolyploidy in the Cercidoideae. If a *Cercis* progenitor contributed one of the subgenomes to the allopolyploid Cercidoideae lineage, then a *Cercis* gene should have greater affinity with one of the two WGD-derived paralogs in, for example, *Bauhinia*. Indeed, this is seen in the consensus species/WGD phylogeny (Fig. 4a).

### Exemplar gene families and utility for studies of evolution of symbiotic nitrogen fixation in the legumes

The consensus topology in Fig. 4a is also seen in individual gene families – as, for example, in Fig. 4c, which happens to be a gene family that has a key role in symbiotic nitrogen fixation (SNF). This family contains MtNSP2, named for the *M. truncatula* “Nodulation Signaling Pathway 2” gene.

The other two gene families shown in Fig. 4 deviate from the consensus topology, but in interesting ways. Fig. 4d contains a gene critical for SNF, identified and named independently in studies using *Lotus japonicus* and *Medicago trunctula*: LjSYMRK/MtDMI2 (SYMbiotic Receptor Kinase / Doesn’t Make Infections 2). In this family, orthologs are present in two nodulating species in the Caesalpinioideae, *C. fasciculata* and *A. crassicarpa*; but not in the (non-nodulating) *S. tora*. Significance of this presence-absence pattern needs to be tempered, however, by the existence of orthologs from the other non-nodulating species in the family: *Bauhinia, Phanera, Cercis*, and *Sindora*.

Fig. 4e contains LjNIN (Nodule Inception), which has been shown to be critical to SNF (Schauser *et al*., 1999; Marsh *et al*., 2007; Griesmann *et al*., 2018). In this gene family, the only legume species present in the family are those that nodulate; and all other non-nodulating species included in this study are absent. Specifically, the nodulating species present in this gene family are *L. japonicus, M. truncatula, P. vulgaris, A. evenia* (Papilionoideae) and *A. crassicarpa* and *C. fasciculata* (Caesalpinioideae). The non-nodulating species that are all absent from this gene family are *S. glauca* (Detarioideae) and *C. canadensis*, *P. championii*, and *B. variegata* (Cercidoideae).

### Ks and phylogenomic analyses indicate independent WGD events in at least four legume subfamilies

Analyses of silent-site mutations between species pairs can be used to examine relative evolutionary rates associated with both speciations and WGDs. The plots in Fig. 5 show Ks peaks for selected species pairs. The resolution in these plots is higher than in many Ks analyses because the Ks values are taken from the modal Ks value per synteny block in the indicated species comparison, rather than from individual gene pairs.

**Fig. 5.**
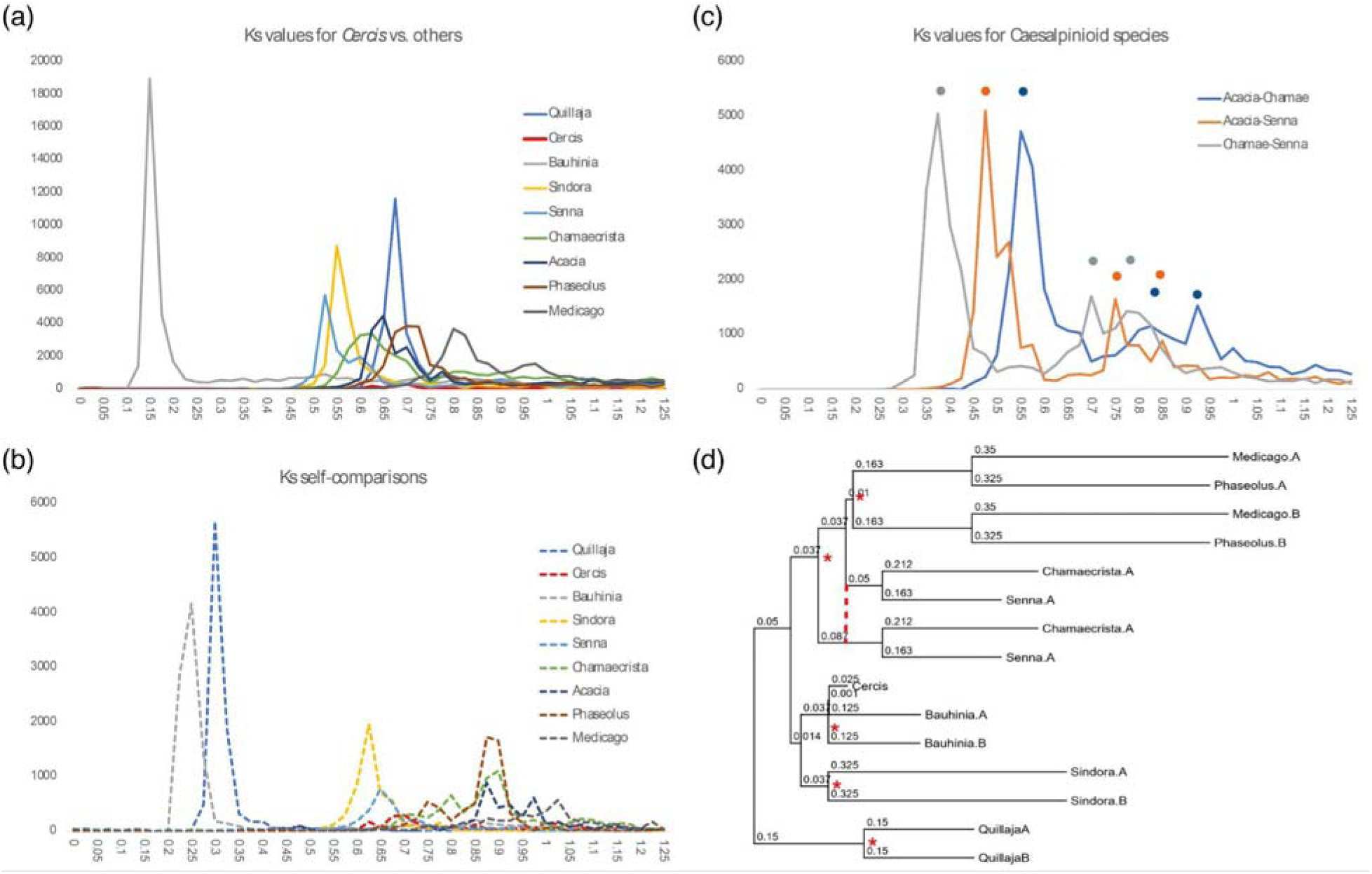
Ks distributions and phylogeny with Ks-derived branch lengths. In panels (a) and (b), modal Ks values were calculated for synteny blocks based on gene-pair matches, and those modal values used in place of the individual gene-pair values. (a) Comparisons between *Cercis* and other species (peaks representing speciations). (b) Ks values for species self-comparisons (peaks representing WGD events). (c) Ks peaks for comparisons among species in the Caesalpinioideae: *Acacia crassicarpa, Chamaecrista fasciculata, Senna tora.* Dots show locations of Ks peaks – on the left indicating speciations, and on the right indicating a whole genome duplication. (d) Phylogeny derived from combined gene families (see Fig. 3 for details), with branch lengths determined algebraically from Ks peaks.

In Fig. 5a, showing Ks values between *Cercis* and the indicated species (including *Cercis* compared with itself), the amplitude of *Cercis-Cercis* plot is near zero in the Ks range shown. The peak for the *Cercis-Cercis* comparison is near Ks=2.0 (not shown), consistent with the only duplication in *Cercis* being much older than the origin of the legumes. The Ks peak with the smallest value is with *Bauhinia*, which is consistent with *Cercis* and *Bauhinia* being relatively close sister taxa within the Cercidoideae. All other peaks in Fig. 5a are in the range 0.5-0.85, reflecting the substantial divergence with the other selected species, all of which are in other subfamilies (and another plant family all together in the case of *Quillaja*).

In Fig. 5b, showing Ks peaks for self-comparisons for each species, *Cercis* again is notable in its near-absence (several-fold lower in amplitude than for the other species self-comparisons). The Ks peaks for *Bauhinia* and *Quillaja* are both in small Ks bins (0.25 and 0.325 respectively), reflecting the relatively recent WGDs in those two taxa (WGDs that must be independent, since they are in different families).

In Fig. 5c, showing comparisons among the three Caesalpinioideae species included in this study, there is a strong primary peak reflecting speciations (*Acacia-Chamaecrista*, *Acacia-Senna*, and *Chamaecrista-Senna*); and an intriguing doubled (bimodal) peak in each comparison at more distant (older) Ks bins: at 0.7 and 0.8 for *Chamaecrista-Senna*, 0.75 and 0.85 for *Acacia-Senna*, and 0.825 and 0.925 for *Acacia-Chamaecrista*. For each species pair, these secondary double peaks are separated by 0.1 Ks units. A secondary peak is expected to represent a WGD near the base of the Caesalpinioideae; but a doubled secondary peak may represent alternate evolutionary paths associated with allopolyploidy, as depicted in the schematic in Fig. 4b.

Because rates of silent-site mutations may differ in different lineages, the values need to be considered in a phylogenetic context. Projecting the Ks values from the species pairs (Table S3) onto the consensus topology from Fig. 4a and resolving the branch lengths algebraically gives the approximate branch lengths in Ks units. Both the phylogenetic and Ks analyses support independent WGDs in each of the examined subfamilies. Some lineages have apparently been evolving much faster than others (by the metric of silent-site mutations), with *Medicago* having accumulated changes at nearly twice the pace of *Cercis* since their divergence from the common legume ancestor (Ks distances to the common ancestor of ~0.5 for *Medicago* and 0.225 for *Cercis*).

### Gene expansion and contraction by comparing to representative species

Analyses of gene expansion and contraction at the level of subfamily showed the largest number of changes in Papilionoideae (represented by *Medicago* and *Phaseolus*), with an increase of 632 gene families and a decrease of 592 (Fig. 6a). At the species level, *Bauhinia* (Cercidoideae), showed the highest number of gene family changes, with 5,243 increases and 728 decreases; while *Cercis* had the lowest number of gene families, with 789 increases and 3,654 decreases. This is consistent with other indications of slower evolution in *Cercis* than other legumes – including *Bauhinia* within the same subfamily.

**Fig. 6.**
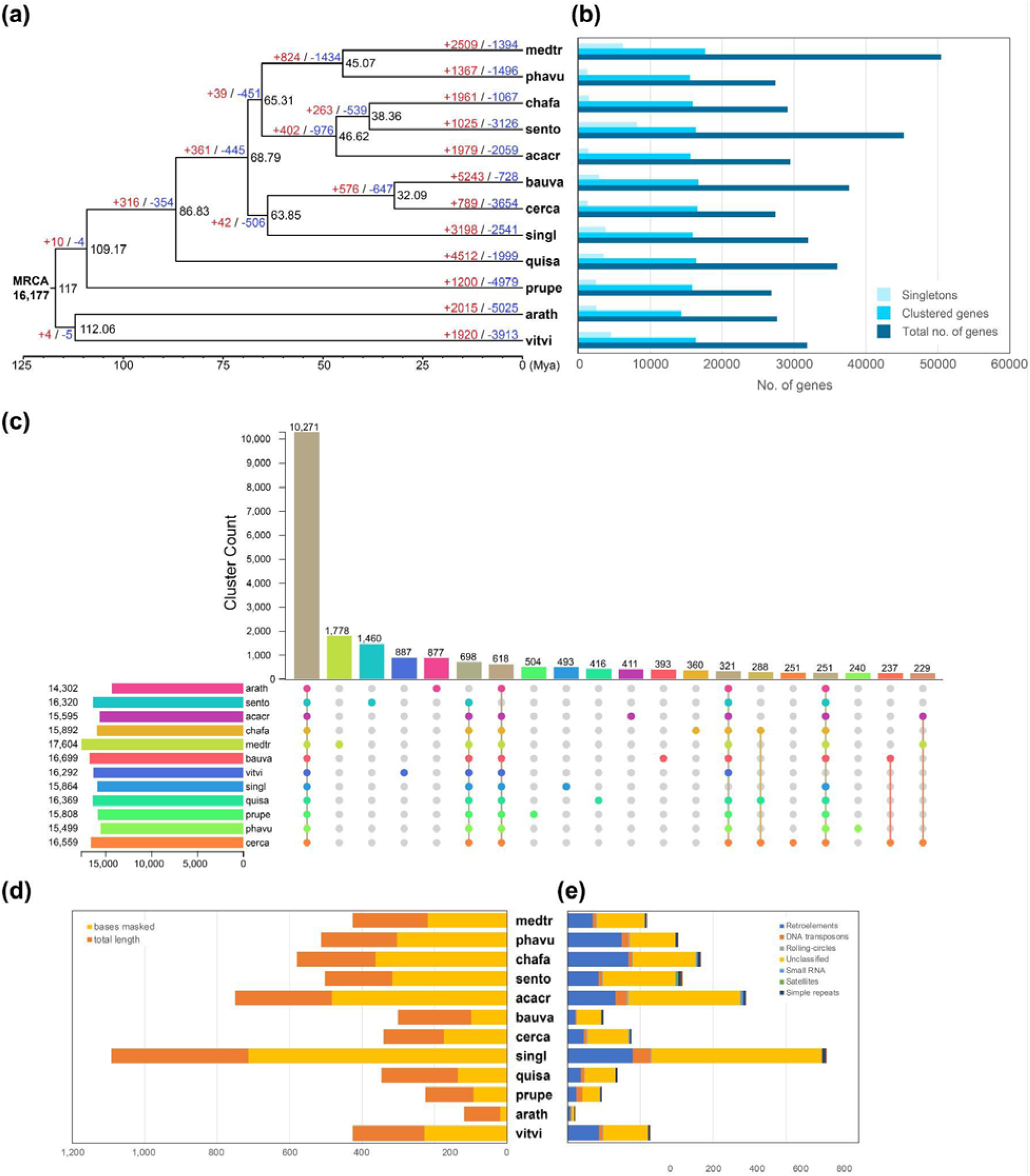
Genomic comparison of *C. canadensis* and *C. fasciculata* with related species. (a) Phylogenetic tree with related species. Estimated divergence times are shown on the node, which indicates the number of expansions (red) and contractions (blue) relative to the gene family with the most recent common ancestor (MRCA). The divergence time was calculated based on 117 Mya for *Medicago* and *Vitis*. (b) Bar graph comparing the total number of genes, clustered genes (orthogroup), and singleton numbers for the species used in the phylogenetic analysis. (c) Upset plot of clustered genes. The number clustered in all species is 10,271. (d) Bar graph showing the proportion of repetitive sequences to genome size. (e) Cumulative bar graph showing the proportion of repetitive sequence content for each species.

In the GO enrichment analysis of these increased and decreased gene families, *Cercis* was compared to *Bauhinia*, and *Chamaecrista* was compared to *Senna*. When comparing *Cercis* and *Bauhinia*, there was an enrichment of terms for defense response, signal transduction, and pollen recognition. Although both species are bisexual flowers, the fact that signal transduction and pollen recognition are most expanded in *Cercis* suggests that they may have unique recognition systems (Figs S3, S5).

In comparison to *Chamecrista* and *Senna*, the top 30 expanded clusters are significantly enriched in signal transduction, pollen recognition, and auxin-activated signaling pathway functions (Figs S4). Interestingly, the auxin-activated signaling pathway is consistent with what we expected since *Chamaecrista* produces nodules unlike *Senna*. It is expected that the flavonoids produced by *Chamaecrista* act as auxin transport inhibitors during the pre-nodule infection stage, affecting the promotion of nodule primordial cell division at the nodule site (Figs S4, S6). Orthologous cluster analysis revealed that 10,271 clusters were common to all species, 251 clusters were unique to *Cercis* and 360 clusters were unique to *Chamaecrista* (Fig. 6c).

### Divergence time estimation

Based on a divergence time of 117 Mya between *Medicago* and *Vitis* (Wikström *et al*., 2001), we estimated the divergence of the other lineages based on phylogenetic topology and relative branch lengths. The following four legume subfamilies diverged at 68.8 Mya. This is similar to the radiometrically dated Cretaceous-Paleozoic (K-Pg) boundary of 66 Mya (Renne *et al*., 2013). In our calculations, the estimated divergence time of the Papilionoideae and Caesalpinioideae was at 65.3 Mya, followed by Detarioideae and Cercidoideae at 63.9 Mya. *Cercis* and *Bauhinia* (in the same subfamily) diverged at 32.1 Mya (Fig. 6a).

### Centromeric arrays in *Cercis* and *Chamaecrista* suggest divergent evolutionary histories

Tandem repeats in *C. canadensis* and *C. fasciculata* were identified using ULTRA (Olson & Wheeler, 2018), The most abundant repeats with length greater than 90 bases were evaluated for chromosomal position and array size. Based on these characteristics, putative centromeric repeats were selected and clustered in order to identify consensus centromeric repeat sequences (Data S1).

The arrays of probable centromeric repeats are strikingly large in *C. canadensis*, extending up to 12 Mb and averaging 6 Mb. In total, they comprise approximately 12.6% of the total genome size. In contrast, the largest centromeric arrays in *C. fasciculata* comprise approximately 6.2% of the total genome size. The centromeric arrays generally occur in different (non-syntenic) locations – the arrays in *Cercis* disrupting synteny with *Chamaecrista* and vice versa (Figs S7-8). The pattern of synteny disruption suggests that centromeric arrays have originated (or moved) subsequent to the divergence of the respective lineages.

## Discussion

The taxa selected as the focus of this work were chosen in order to help answer questions about the early diversification and evolution of the legume family, and to provide resources for further study of the evolution of symbiotic nitrogen fixation.

*Cercis*, as the earliest diverging lineage in the Cercidoideae (which itself is early-diverging within the legumes), is of particular interest due to its lack of WGD, its slow evolutionary rate, and its similarity to the inferred chromosome structure in the legume progenitor. As the only legume established to this point to be without a WGD in the span of legume evolution, *Cercis* offers a unique model for the study of evolution of this large and diverse family.

Synteny, phylogenomic, and Ks analyses confirm that *C. canadensis* does not have a recent WGD within the timeframe of the legume family. Limited evidence of older duplications are evident, consistent with the ~135 Mya gamma triplication (Jiao *et al*., 2012). In addition, independent WGD events are evident in the other four legume subfamilies that we examined in this study: the Papilionoideae, Caesalpinioideae, Detarioideae, and Cercidioideae. The estimated divergence time of each subfamily in the 63.9-68.8 Mya (Fig. 6a).

We also infer a probable karyotype of seven chromosomes in the legume ancestor, structurally similar to the current *Cercis* chromosomes, albeit with some rearrangements. Although some other examined species are found to have complex rearrangements (particularly in the Papilionoideae), the others are generally well approximated as a doubling of the *Cercis* genome, followed by a small number of splits, fusions, and inversions, which can be crucial for reconstructing the genomic history of the legume family.

We speculate that the unusually large centromeric arrays in *C. canadensis* (comprising roughly 12% of the genome sequence), may be related to the stability of the chromosome structure over the ~70 million year history of legume evolution - as both cause and effect. In particular, centromeric arrays may tend to grow if undisturbed by rearrangements; and large centromeres may also aid in maintaining chromosome structure by providing “unmissable” mitotic attachment points.

Analysis of the expansion and contraction of duplicated genes and gene families confirms that the retention of duplicated genes has not been random, with some families generally retaining post-WGD duplicates (including in lineages with independent WGDs), and other families tending to fall back to single-copy status. Stochastic gene loss may have been notably important regarding SNF, where the pattern of presence and absence of this trait across the nitrogen-fixing clade has been modeled as a small number of independent gains and numerous losses (Griesmann *et al*., 2018; Kates *et al*., 2024).

*Chamaecrista* is of particular interest due to its capacity for SNF -- in contrast with many other lineages in the Caesalpinioideae subfamily -- including genera such as *Senna*, which is a sister genus within the Cassiinae tribe (and within the broader Mimosoideae–Caesalpinieae–Cassiinae (MCC) clade in the Caesalpinioideae). *Chamaecrista* has long been proposed as a model for examination of SNF within the Caesalpinioideae, due in part to characteristics that make *Chamaecrista* suitable for experimentation (Singer *et al*., 2009). In particular, *C. fasciculata* is physically small, with short generation time, it can be outcrossed or selfed, and it exhibits considerable diversity across the North American habitats in which it is found.

The genome assemblies and annotations, together with other resources representing the four largest subfamilies in the legumes, permit construction of gene families that robustly capture the core genic complement of the family. These gene families support allopolyploid genome duplication events in both the Cercidoideae and Caesalpinioideae. This result helps explain why it has been so difficult to resolve the backbone topology for the legume phylogeny. Considerable discordance is seen in the placement of genes from Caesalpinioideae, such that WGD-derived paralogs from species such as *Chamaecrista* often do not resolve as sister to one another, but rather as alternately sister to genes from other subfamilies. An allopolyploid model is consistent with this observed pattern of discordance in gene families for Caesalpinioid sequences.

An important general corollary of allopolyploidy within a taxonomy is that it may not be possible to faithfully represent the history of species relationships with a bifurcating phylogenetic model. Rather, a reticulate model is needed. Furthermore, particular genes may have followed different evolutionary histories due to effects such as incomplete lineage sorting, gene conversion, and segmental or chromosomal replacement following allopolyploidy.

Although the work presented here did not focus primarily on nodulation, the availability of a high-quality genome assembly and annotations for *C. fasciculata* are expected to be of use in such studies in the future. Examination of key nodulation-related gene families such as SYMRK and NIN show the utility of high-quality genic sequence from *Chamaecrista* and *Senna* -- the SYMRK gene family showing retention of both WGD-derived *Chamaecrista* genes but loss of both from *Senna*; and NIN showing presence of orthologs in all examined nodulating species and absence from all non-nodulating species.

Within the Caesalpinioideae, nodulation is present in approximately nine lineages and absent in a comparable number (Sprent *et al*., 2017; Kates *et al*., 2024). Nodulation is present in most genera in the Papilionoideae, and absent in the other four legume subfamilies. The pattern of taxonomically scattered presence and absence of the trait in the Caesalpinoideae has been modeled as due to repeated, scattered losses of SNF (Kates *et al*., 2024). We speculate that this pattern of loss could have been facilitated by the likely allopolyploid history of this subfamily. For example, if one of the progenitor species lacked SNF (either due to loss or non-gain) and the other progenitor had SNF capacity, then the allopolyploid merger might have produced a new polyploid species that had capacity for diversification, but that was also vulnerable to stochastic loss of genes crucial to SNF.

In the SYMRK/DMI2 family (Fig. 4d) (SYMRK identified in *Lotus japonicus* and the ortholog DMI2 identified in *Medicago truncatula*), two WGD-derived paralogs are present in *Acacia* (within the Mimosoid group, where nodulation predominates among most taxa). In *Chamaecrista*, one of two paralogs has evidently been lost. Both paralogs have been lost from (non-nodulating) *Senna*. While we can’t establish the exact timing of the losses or the precise historical functions of these orthologs given data from this project, this pattern of presence and loss is consistent with inheritance of progenitor SYMRK/DMI2 genes from two species that merged to form the early allopolyploid founder of the Caesalpinoideae. In this model, those two genes may already have acquired differing functions. Both were evidently retained in *Acacia* and only one in *Chamaecrista*. Both were lost in *Senna*.

A similar story may apply for the NIN gene family (Fig. 4e). In that case, at least one functioning progenitor gene must have been present prior to the origin and radiation of the Papilionoideae and Caesalpinioideae. One WGD paralog was evidently lost from the *Acacia* (mimosid) lineage, but both WGD-derived paralogs have been retained in each of the Papiloionoideae and in *Chamaecrista*. Intriguingly, one of the *Chamaecrista* genes (ChafaH1.1G219100) resolves sister to the described NIN gene, Medtr5G099060.

## Conclusions

The work here describes the high-quality genomes and annotations for *Cercis canadensis*, eponymous for the *Cercidoideae* legume subfamily; and for *Chamaecrista fasciculata* in the Caesalpinoideae subfamily. These species are well placed taxonomically to aid inferences about key features of legume evolution, including the legume ancestral karyotype and the respective timing of subfamily origins and WGDs early in the legume radiation. Both *Cercis* and *Chamaecrista* show evidence of allopolyploidy – with the progenitor of *Cercis* likely contributing one subgenome to an allopolyploid event that gave rise to the remaining species in the Cercidoideae. In the Caesalpinoideae, the preponderance of gene families and Ks analyses suggest merger of two species that diverged along the taxonomic grade leading to the Papilionoideae, and then merged to give rise to the allopolyploid Caesalpinoideae. Such an allopolyploid merger early in the evolution of SNF may help to explain the very uneven pattern of SNF presence and absence across the diverse Caesalpinoideae. Finally, a finding of allopolyploidy during the origin of the legumes provides an important example of diversification that is not modeled sufficiently with a standard bifurcating phylogeny.

## Supporting information

Supporting Information Methods S1 and Results S2

## Acknowledgements

This work (proposal: 10.46936/10.25585/60001405) conducted by the U.S. Department of Energy Joint Genome Institute (https://ror.org/04xm1d337), a DOE Office of Science User Facility, is supported by the Office of Science of the U.S. Department of Energy operated under Contract No. DE-AC02-05CH11231. The work was also supported by the United States Department of Agriculture, Agricultural Research Service (USDA-ARS) CRIS Project 5030-21000-071-000D. This research used resources provided by the SCINet project and/or the AI Center of Excellence of the USDA Agricultural Research Service, ARS project numbers 0201-88888-003-000D and 0201-88888-002-000D. This research used resources of the National Energy Research Scientific Computing Center, a DOE Office of Science User Facility supported by the Office of Science of the U.S. Department of Energy under Contract No. DE-AC02-05CH11231 using NERSC award BER-ERCAP0027438. The USDA is an equal opportunity provider and employer. Mention of trade names or commercial products in this article is solely for the purpose of providing specific information and does not imply recommendation or endorsement by the U.S. Department of Agriculture.

## Author Contributions

HL and SBC conducted primary analyses, drafted the review, and managed project data. HL, JSS, and SBC conducted gene family, phylogenomic and Ks analyses. SBC and JSS collected plant tissue for genome sequencing. BDJ contributed analysis of genomic repeats. JG, JS, JJ, and RW conducted the genome sequencing and assembly. MW, JW, KK, and JE conducted lab work for genome sequencing and annotation. TB and SS generated genome annotations. DG and KB managed genome assembly and annotation work. QX, TH, RB, AL, DS, and LJ carried out synteny analyses and contributed software for genomic analysis. JL-M and LTL reviewed the analyses and edited the manuscript. SBC and JL-M conceptualized and designed the research. DMG, KB, and JL-M provided project management and funding. All authors approved the final draft.

## Data Availability

BioProject for *Chamaecrista fasciculata* var. ISC494698: https://www.ncbi.nlm.nih.gov/bioproject/PRJNA1137390

BioProject for *Cercis canadensis* ISC453364: https://www.ncbi.nlm.nih.gov/bioproject/PRJNA1137384

Genome assemblies and annotations for *Chamaecrista fasciculata* var. ISC494698 and *Cercis canadensis* ISC453364: https://phytozome-next.jgi.doe.gov

Legume gene families and associated phylogenomic analyses: https://data.legumeinfo.org/LEGUMES/Fabaceae/genefamilies/legume.fam3.VLMQ/

Supporting Information Methods S1 and Results S2 (manuscript supplement)

Supporting Information Figures and Tables S3 (manuscript supplement)

