## Supporting Information Methods S1 and Results S2 for "Legume genome structures and histories inferred from *Cercis canadensis* and *Chamaecrista fasciculata* genomes"

**New Phytologist Supporting Information Article title: Legume genome structures and histories inferred from *Cercis canadensis* and *Chamaecrista fasciculata* genomes**

**Authors** : Hyun-oh Lee, Jacob S Stai, Qiaoji Xu, Thulani Hewavithana, Rabnoor Batra, Alex Liu, Brandon D Jordan, Rachel Walstead, Jerry Jenkins, Melissa Williams, Jenell Webber, Jane Grimwood, John T Lovell, Tomáš Brůna, Shengqiang Shu, Keykhosrow Keymanesh, Joanne Eichenberger, Jeremy Schmutz, David M Goodstein, Kerrie Barry, David Sankoff, Lingling Jin, James H Leebens-Mack, Steven B Cannon

**Article acceptance date:**

**The following Supporting Information is available for this article:**

**Supporting Information Methods S1**

**Genome assembly of *Cercis canadensis* and *Chamaecrista fasciculata***

The main assembly consisted of CCS PACBIO sequences, 83.12x for *Cercis* and 106.1x for *Chamaecrista* (20,744 and 20,503 bp average read size, respectively), assembled using HiFiAsm+Hi-C (Cheng *et al.*, 2021). The resulting sequence was polished using RACON (Vaser *et al.*, 2017). Misjoins in the assembly were identified using Hi-C data. The following misjoins were identified and corrected: one in the *Cercis* HAP1 assembly, one in the *Chamaecrista* HAP1 assembly, and 27 in the *Chamaecrista* HAP2 assembly.

Contigs for both species and haplotypes were then oriented, ordered, and joined into chromosomes using the JUICER pipeline (Durand *et al.*, 2016), , and chromosome scale scaffolding was performed with 3D-DNA (Dudchenko *et al.*, 2017). Contigs containing significant telomeric sequence were properly oriented in the assembly. For *Cercis*, a total of 8 contig joins were made to form the final assembly consisting of 7 chromosomes. For *Chamaecrista*, a total of 82 joins were applied to the HAP1 assembly, and 98 joins for the HAP2 assembly. A total of 99.46% of the assembled sequence is contained in the chromosomes for *Cercis* HAP1. For *Chamaecrista* a total of 99.58% and 99.40% of assembled bases integrated into the respective HAP1 and HAP2 chromosomes (Figs S9-10). For each assembly, chromosomes were numbered to be consistent with the first haplotype, and the resulting sequence was screened for retained vector and/or contaminants.

Heterozygous snp/indel phasing errors were identified and corrected as follows. For the *Cercis* assemblies, the aligned 83.12x CCS data identified 84 heterozygous single-nucleotide polymorphisms (SNPs) / insertion and deletions (INDELs)d. Additionally, homozygous SNPs and INDELs were corrected in the release sequence using ~49.7x of Illumina reads (2x150, 400bp insert). For the *Chamaecrista* assemblies, homozygous SNPs and INDELs were corrected in the HAP1 and HAP2 releases using ~61.4x of Illumina reads (2x150, 400bp insert) by aligning the reads using bwa mem (Li, 2013) and identifying homozygous SNPs and INDELs with the GATK’s UnifiedGenotyper tool (McKenna *et al.*, 2010). A total of 308 homozygous SNPs and 7,815 homozygous INDELs were corrected in the HAP1 release, while a total of 333 homozygous SNPs and 7,555 homozygous INDELs were corrected in the HAP2 release.

**Genome annotation of *Cercis canadensis* and *Chamaecrista fasciculata***

Transcript assemblies for *C. canadensis* (both haplotypes) and *C. fasciculata* (both haplotypes) were made from ~1.6B and ~748M pairs of 2X150 bp stranded paired-end Illumina RNA-seq reads, respectively, using PERTRAN, which conducts genome-guided transcriptome short read assembly via GSNAP (Wu & Nacu, 2010) and builds splice alignment graphs after alignment validation, realignment, and correction. To obtain ~476K (for *C. canadensis* HAP1*)*, ~475K (*C. canadensis* HAP2*)*, ~503K (*C. fasciculata* HAP1*)*, and ~502K (*C. fasciculata* HAP2*)* putative full-length transcripts, about 13M (*C. canadensis*) and 8.5M (*C. fasciculata*) PacBio Iso-Seq Circular Consensus Sequences (CCS) were corrected and collapsed by a genome-guided correction pipeline. The pipeline aligns CCS reads to the genome with GMAP (Wu & Watanabe, 2005), corrects small indels in splice junctions, and clusters alignments when all introns are the same or >= 95% overlap for single-exon alignments. Subsequently, PASA (Haas *et al.*, 2008) was used to construct 411,027 (*C. canadensis* HAP1), 409,048 (*C. canadensis* HAP2), 267,332 (*C. fasciculata* HAP1), and 266,538 (*C. fasciculata* HAP2) transcript assemblies by combining the respective transcript assembly sets described above.

A repeat library for each genome was created from *de novo* repeats predicted by RepeatModeler2 (Flynn *et al.*, 2020) on the HAP1 of each species. The predicted repeats underwent functional analysis through InterProScan (Jones *et al.*, 2014), incorporating the Pfam (Mistry *et al.*, 2021) and PANTHER (Mi *et al.*, 2019) databases. Any repeats that displayed significant hits to protein-coding domains were subsequently excluded from each repeat library. Finally, the constructed species-specific repeat libraries were used to soft-mask the respective genomes with RepeatMasker (Smit *et al.*, 2015).

Putative gene loci in each genome were determined by transcript assembly alignments and/or EXONERATE (Slater & Birney, 2005) alignments of proteins from *Arabidopsis thaliana, Glycine max, Populus trichocarpa, Oryza sativa, Solanum lycopersicum, Vitis vinifera, Prosopis alba, Prunus persica*^[1]^*, Aquilegia coerulea*^1^*, Amborella trichopoda*^1^*, Numphaea colorata*^1^*, Cercis canadensis*^[2]^*, Lupinus albus*^2^*, Mimulus guttatus*^2^*, Liriodendron tulipifera*^2^*, Carya illinoinensis*^2^*, Lens ervoides*^2^*, Lotus japonicus*^2^*, Arachis duranensis*^2^*, Medicago truncatula*^2^*, Cicer arietinum*^2^*, Abrus precatorius*^2^*, Cucumis sativus*^2^*, Beta vulgaris*^2^*, Sorghum bicolor*^2^*,* and Swiss-Prot eukaryotic proteome release 2020_06*^1^*, and 2022_04*^2^* to repeat-repeat-masked assemblies. The putative loci were extended up to 2kb in both directions of alignments unless they extended into another locus on the same strand. Gene models in each locus were predicted by homology-based predictors, FGENESH+ (Salamov & Solovyev, 2000), FGENESH_EST (similar to FGENESH+, but using EST to compute splice site and intron input instead of protein/translated ORF), EXONERATE, PASA assembly ORFs (in-house homology constrained ORF finder), and AUGUSTUS trained on the high confidence PASA assembly ORFs and with intron hints from short read alignments. The best-scored predictions for each locus were selected using multiple positive factors including EST and protein support, and one negative factor: overlap with repeats. The selected gene predictions were improved by PASA. The improvement included adding UTRs, splicing correction, and adding alternative transcripts.

PASA-improved gene model proteins were subject to protein homology analysis to the above-mentioned proteomes to obtain a Cscore and protein coverage. Cscore is a protein BLASTP score ratio to the mutual best hit BLASTP score and protein coverage is the percentage of protein aligned to the best of homologs. PASA-improved transcripts were selected based on Cscore, protein coverage, EST coverage, and their CDS overlap with repeats. The transcripts were selected if their Cscore and protein coverage were >= 0.5 or if covered by ESTs. For gene models whose CDS were overlapped by repeats by more than 20%, their Cscore had to be at least 0.9 and homology coverage at least 70% to be selected. The selected gene models were subject to Pfam analysis and gene models without strong transcriptome and homology support whose proteins were more than 30% overlapped by Pfam TE domains were removed. Incomplete gene models, low homology supported without fully transcriptome-supported gene models, short single exon (< 300 bp CDS) without protein domains nor good expression, and repetitive gene models without strong homology support were manually filtered out.

**Supporting Information Results S2 - Analysis of Repetitive Sequences and Gene Family Composition**

**Genome composition of repetitive sequences**

By determining the proportion of repetitive sequences in each genome, we estimate the extent to which repetitive sequences have affected genome size. With the exception of the *S. glabra* genome, which is 1.09 Gb, the median genome size is 426 Mbp. *C. canadensis* was approximately 86 Mbp smaller than the median genome size, while *C. fasciculata* was 154 Mbp larger than median value (Fig. 6d).

The repetitive sequences influence various genome sizes. The proportion of repetitive sequences in the genomes was lowest in *A. thaliana* with 16.1% (of 19.2 Mbp) and highest in *S. glabra* with 65.4% (of 714 Mbp). On median value, these genomes contained 52.3% repetitive sequences. While *S. glabra* has the largest repetitive sequence size in absolute, there are a few species that stand out when looking at the proportions in each genome.

Comparing the average values by repeat class category, unclassified repeats accounted for the highest proportion of 41.2-68.5% in all species, followed by retroelements of 21.1-49.1%, DNA transposons of 2.4-18.3%, simple repeats, rolling circles, low complexity, small RNA, and satellites. Evaluating retroelement composition, *P. vulgaris* and *C. fasciculata* show expansion of retroelements relative to the other compared species, accounting for 49.1 and 46.1%, respectively. These proportions are more than twice the rate of *B. variegata*, which has a rate of about 21.1%. Next, for *P. persica*, the rate of DNA transposons is 18.3%, which is about 7.6 times higher than *C. fasciculata*, which is 2.4%. The species-specific distribution of these specific repeats in the genome is expected to have had a significant impact on evolution (Fig. 6e;Table S2).

In particular, a comparison of Cercidoideae, *C. canadensis* and *B. variegata* shows that the genome sizes are similar; 340 (*C. canadensis)* vs. 300 Mbp (*B. variegata*). However, the repetitive sequences increased about 1.8-fold (174/98 Mbp). Specifically, rolling circles and small RNA increased by 4.2-fold, DNA transposons by 2.4-fold, retroelements by 2.1-fold, and unclassified by 1.7-fold, whereas simple repeat and low complex decreased by 0.7-fold and 0.8-fold, respectively (Fig. 6e;Table S2).

Regions with higher densities of these repeat sequences correlate with lower gene densities (Fig. 1**)**. The repeat density maps created by repeat class also showed where the repeats were distributed on the chromosome. Interestingly, *Cercis* tended to have a distinctly higher or lower density of repetitive sequences compared to *Chamaecrista*. In *Cercis*, with the exception of chromosome 6, there were large blocks where no repeats were present. In addition, a region with a very high density of unknown repeats, presumably the centromere, was strongly identified ( Fig. S7). Whereas, in *Chamaecrista*, these regions were identified on only chromosomes 1, 2, 3, and 5 (Fig. S8).

**Retention of duplicated genes is not random.**

Looking at gene families in which WGD-derived gene pairs are retained, a core of around 500-800 gene families are consistently retained in species from different legume subfamilies. For example, between *Phaseolus, Senna,* and *Sindora* (from the Papilionoideae, Caesalpinioideae, and Detarioideae subfamilies), 734 gene families are in common, among gene families with retained duplicates for those species. Adding *Medicago* and *Bauhinia* (from the Papilionoideae and Cercidoideae subfamilies), the intersection drops only moderately, to 466. Only when adding *Cercis*, which lacks a recent WGD, does the intersection drop substantially - to 21 gene families with gene duplicates across all included species.

If reversion to singletons were random and independent in each subfamily, then the number of gene families with retained duplicates in species from three subfamilies would be around 160 rather than the observed ~500-800. (The expected intersection is calculated as: (2200 retained duplicates per species / 8122 low-copy, conserved families) raised to the third power, times 8122 gene families; or: 0.27^3 x 8122 = 160).

Among the 466 genes retained across all four subfamilies, there is enrichment of genes involved in DNA-binding transcription factor, phosphatidylinositol phosphate kinase activity, and chromatin binding (molecular function ontology categories); and for regulation of transcription, phosphatidyl inositol metabolic process, and cytoskeletal organization (biological process ontology categories).

**Following legume WGDs, most gene families have reverted to single-copy.**

Considering gene families that are represented in all six examined legume species and at low-copy (one or two members in each species), a large proportion of the families have reverted to single-copy status. In each of *Senna*, *Medicago*, and *Phaseolus*, 73% are single-copy, and 62% are single-copy in *Sindora*. *Bauhinia* has 40% at single-copy, consistent with its more recent duplication. *Cercis* is a marked exception, with 95% being single-copy – reflecting its lack of a recent whole-genome duplication.

***Cercis* is well represented across legume gene families**

In gene families calculated in this study, including six legume representatives from four subfamilies and four non-legume outgroup species, *Cercis* genes are present in 17924/24186 = 74.1% of the gene families. Of the 20080 gene families with at least two legume species represented, 17748 (88.4%) of these families include at least one *Cercis* gene.

1. Proteomes aligned only to *C. canadensis*
2. Proteomes aligned only to *C. fasciculata*
