## Supporting Information Figures and Tables S3 for "Legume genome structures and histories inferred from *Cercis canadensis* and *Chamaecrista fasciculata* genomes"

**New Phytologist Supporting Information Article title: Legume genome structures and histories inferred from *Cercis canadensis* and *Chamaecrista fasciculata* genomes**

**Authors** : Hyun-oh Lee, Jacob S Stai, Qiaoji Xu, Thulani Hewavithana, Rabnoor Batra, Alex Liu, Brandon D Jordan, Rachel Walstead, Jerry Jenkins, Melissa Williams, Jenell Webber, Jane Grimwood, John T Lovell, Tomáš Brůna, Shengqiang Shu, Keykhosrow Keymanesh, Joanne Eichenberger, Jeremy Schmutz, David M Goodstein, Kerrie Barry, David Sankoff, Lingling Jin, James H Leebens-Mack, Steven B Cannon

**Article acceptance date:**

**The following Supporting Information is available for this article:**

**Supporting Information Figures and Tables S3**

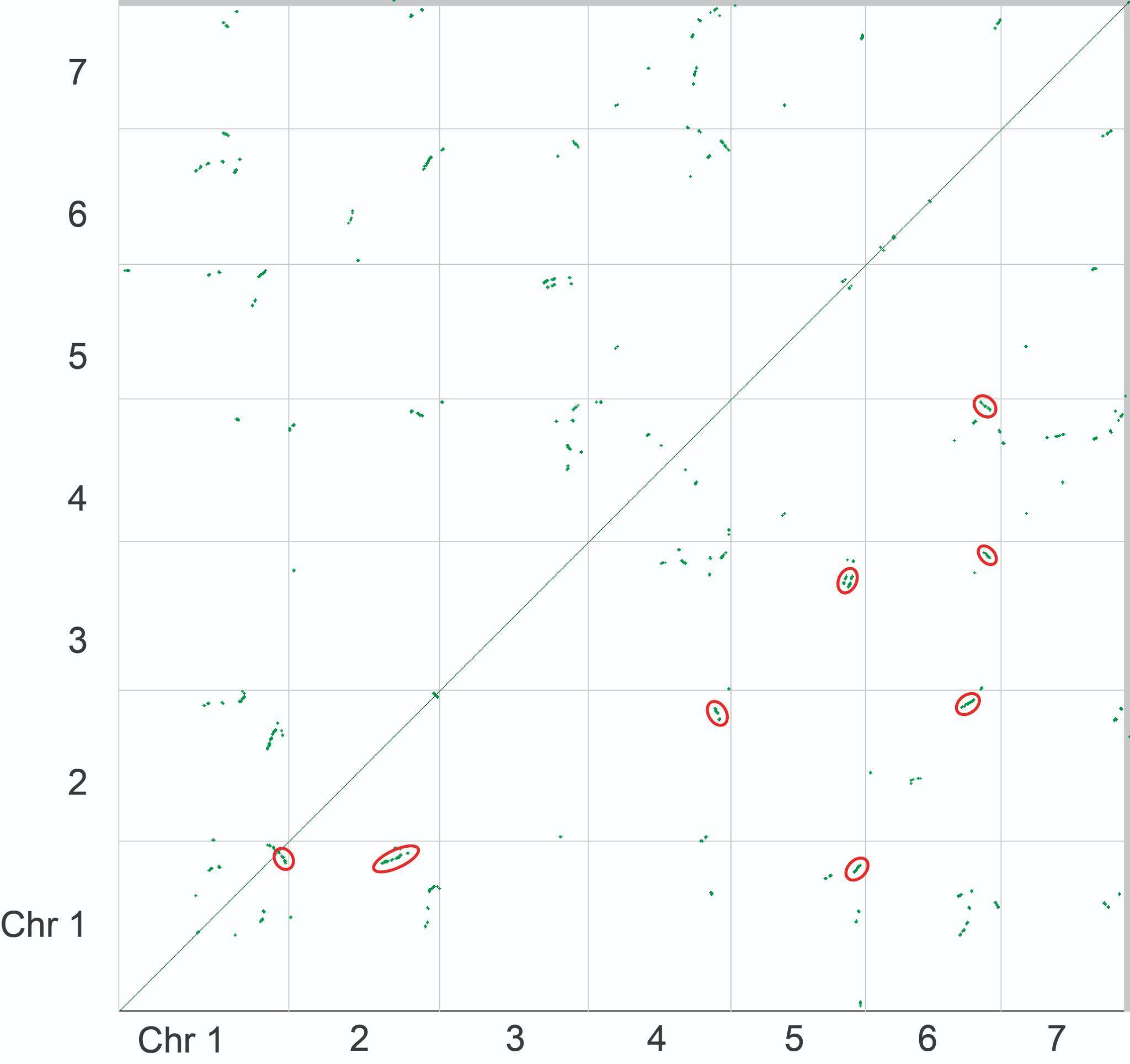

**Supplementary Figure 1.** Syntenic dotplots of the *C. canadensis* genome, red circles indicate traces of old (>100 Mya) duplication(s).

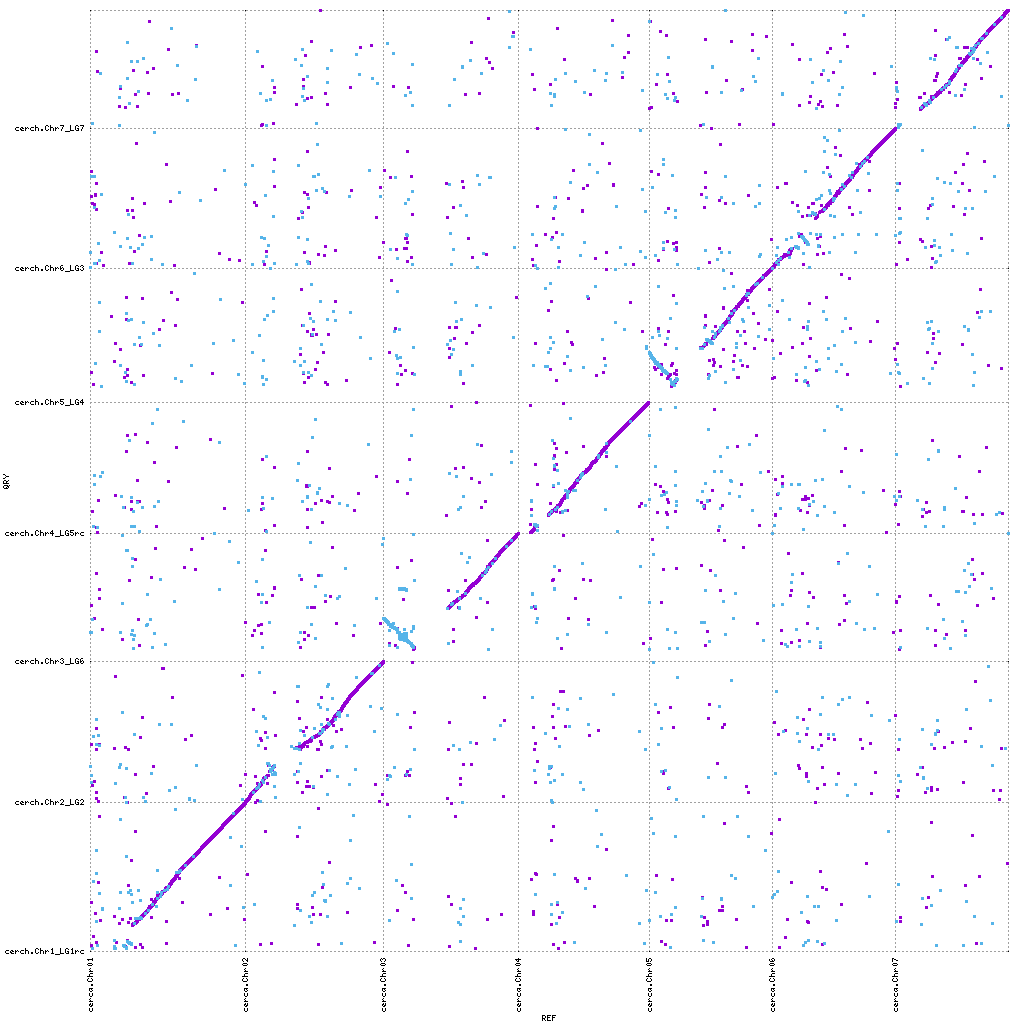

**Supplementary Figure 2.** Syntenic dotplots of the *C. canadensis* (x-axis) vs. *C. chinensis* (y-axis)*.* There are some mismatched regions, and inversions have occurred on chromosomes 3 and 5.

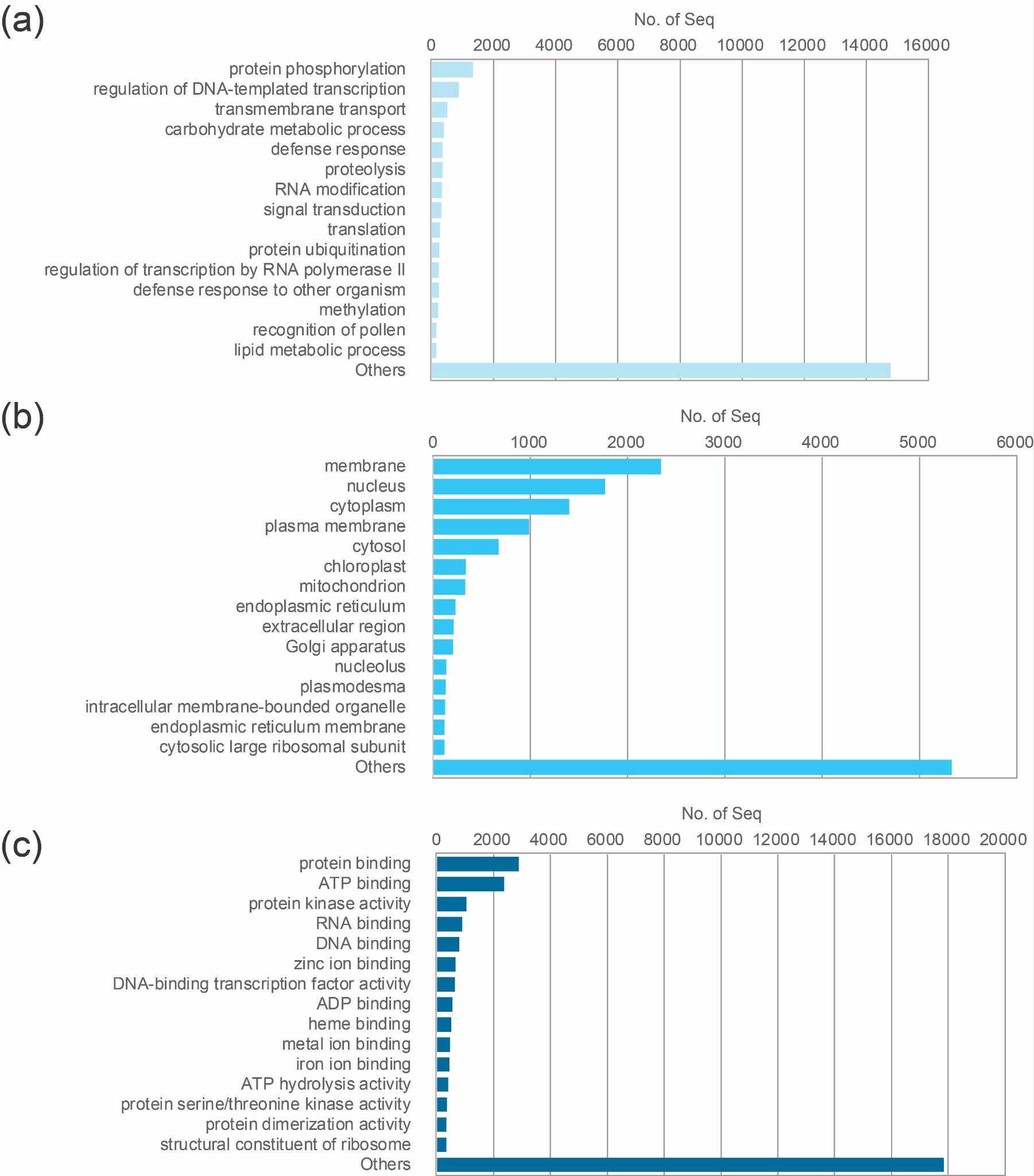

**Supplementary Figure 3**. Gene ontology (GO) enrichment analysis of *C. canadensis* by three functional groups. (A) biological processes (B) cellular components (C) molecular function

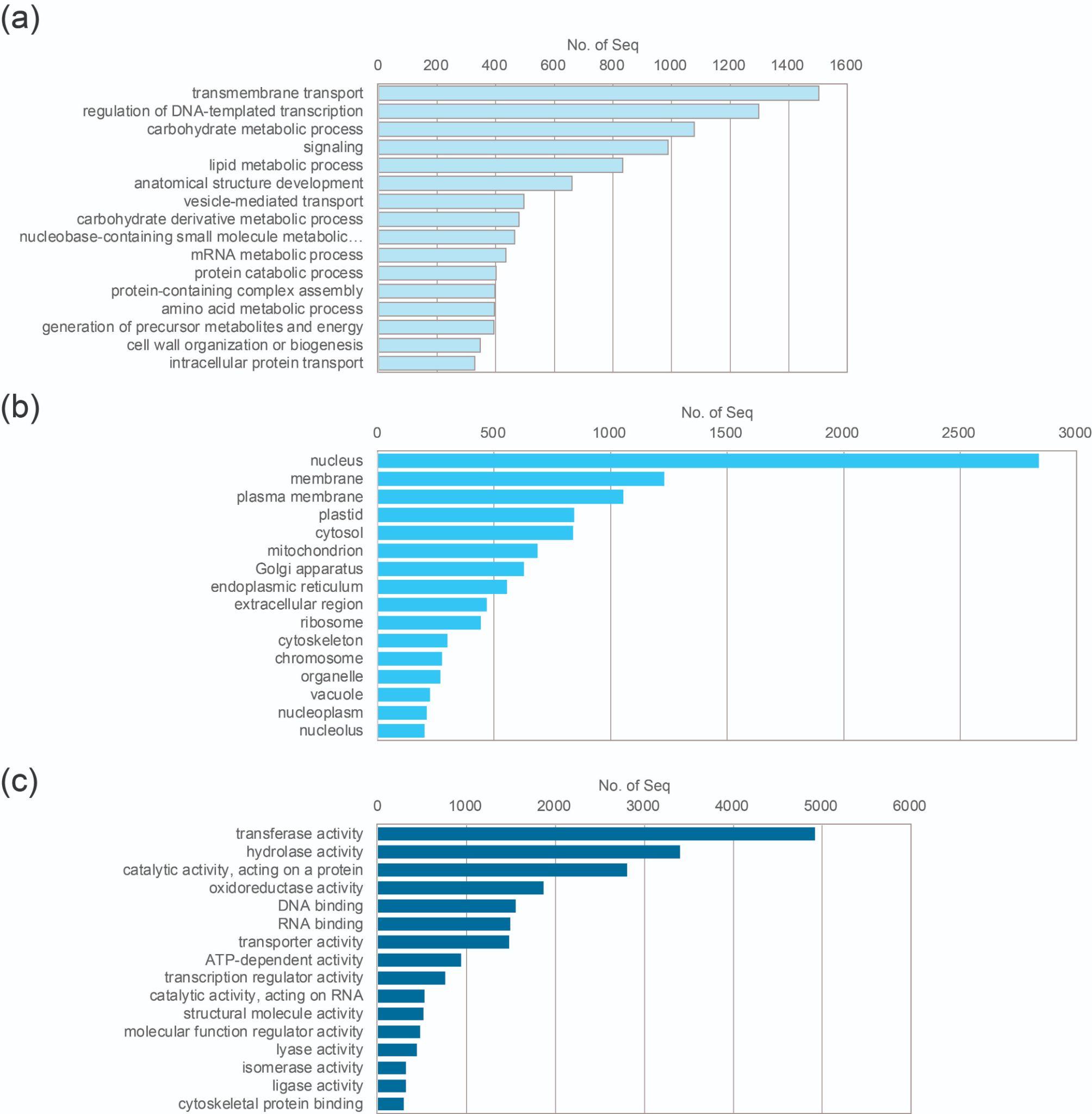

**Supplementary Figure 4**. Gene ontology (GO) enrichment analysis of *C. fasciculata* by three functional groups. (A) biological processes (B) cellular components (C) molecular function

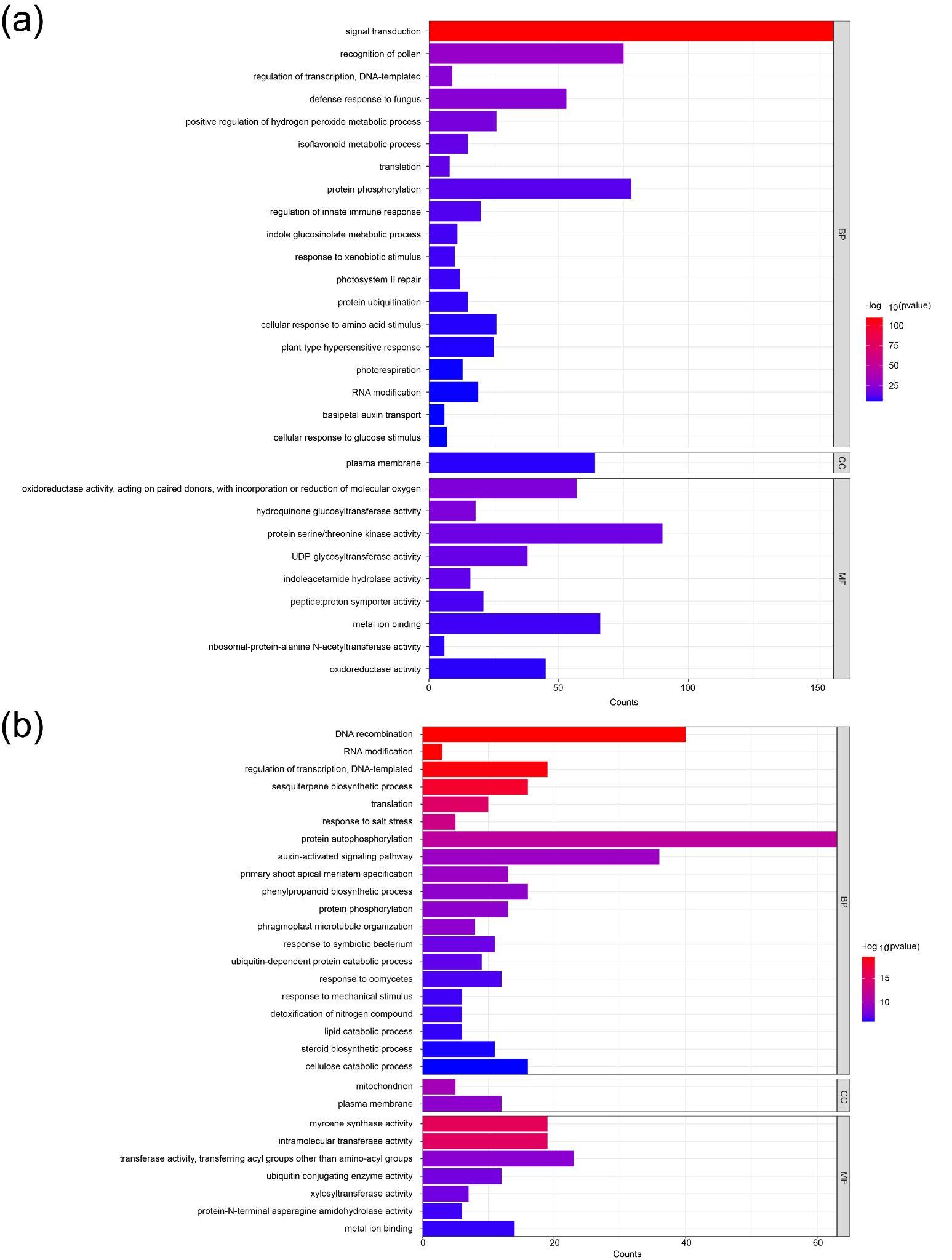

**Supplementary Figure 5.** GO enrichment analysis in the comparison between *C. canadensis* and *B. variegata*. The top 30 terms with p-value of -log10 were shown. The ontology categories are divided into three; BP: biological process, CC: cellular component, MF: molecular function. (A) represents the increased GO terms (B) represents the decreased GO terms.

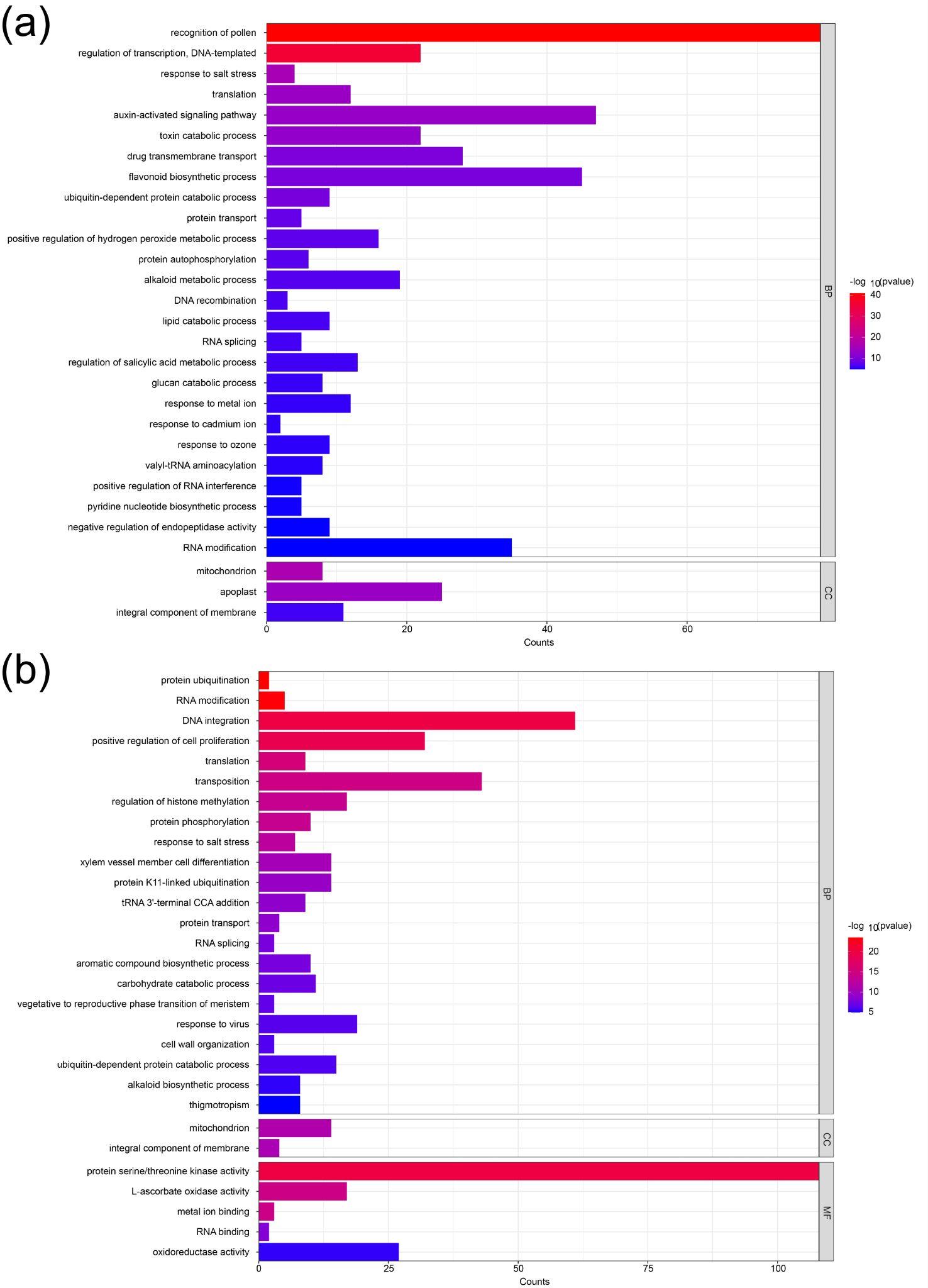

**Supplementary Figure 6.** GO enrichment analysis in the comparison between *C. fasciculata* and *S. tora*. The top 30 terms with p-value of -log10 were shown. The ontology categories are divided into three; BP: biological process, CC: cellular component, MF: molecular function. (A) represents the increased GO terms (B) represents the decreased GO terms.

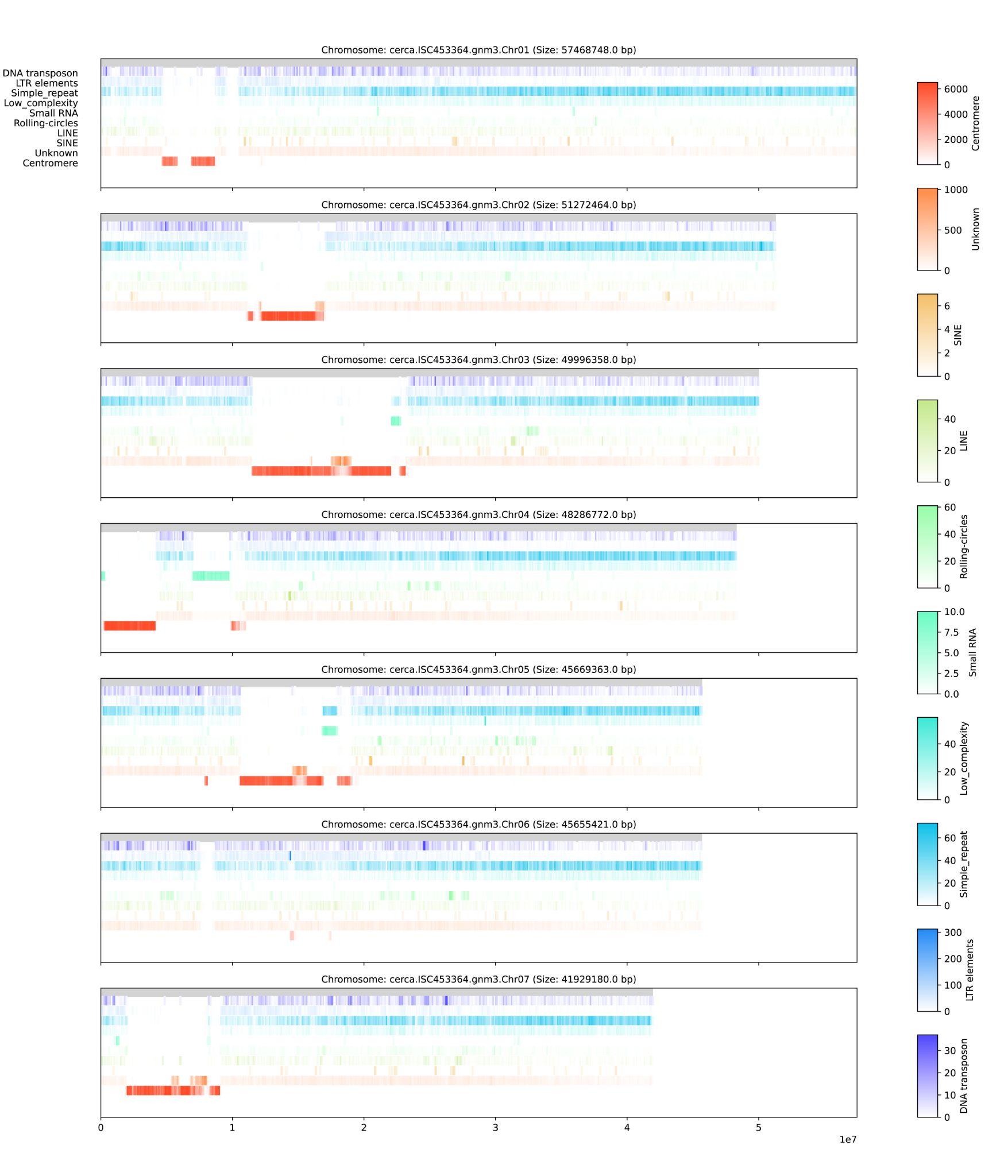

**Supplementary Figure 7.** Density diagram by repeat class in *C. canadensis*. Since density is different for each repeat class, it is shown separately on the right legends.

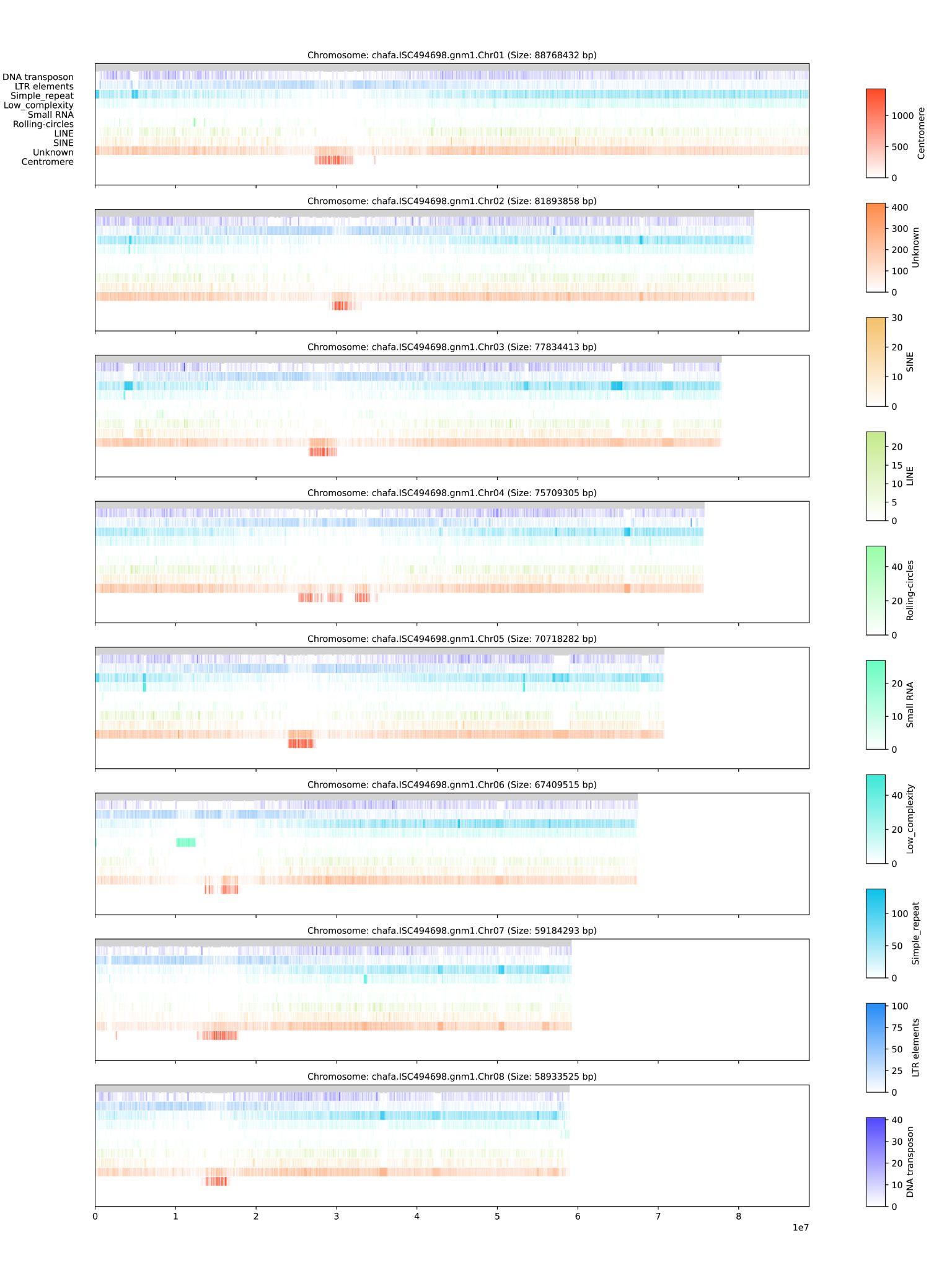

**Supplementary Figure 8.** Density diagram by repeat class in *C. fasciculata*. Since density is different for each repeat class, it is shown separately on the right legends.

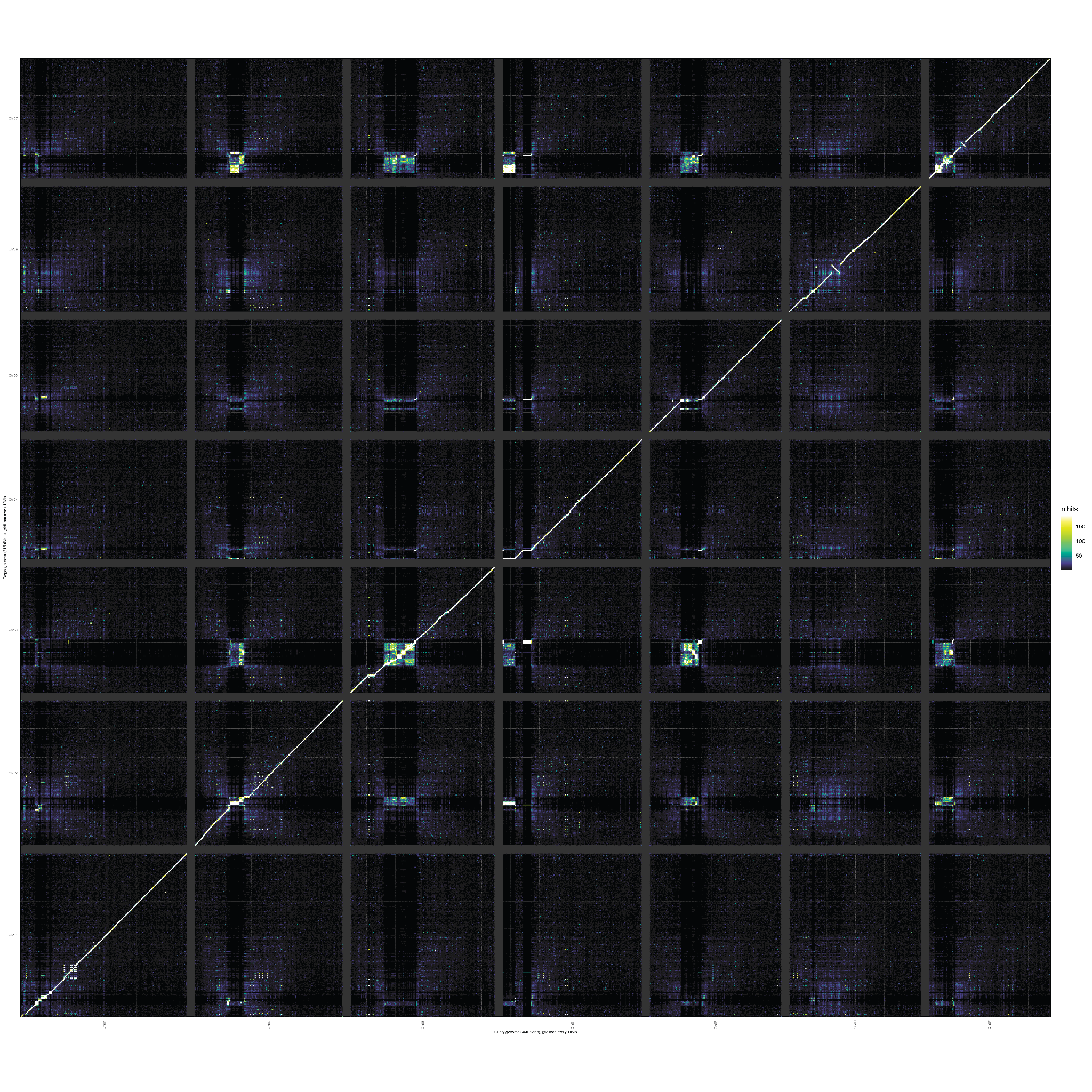

**Supplementary Figure 9.** Comparisons of haplotype assemblies for *C. canadensis*. Regions of higher densities (lighter) show higher identities, indicating either conserved synteny or repetitive sequence.

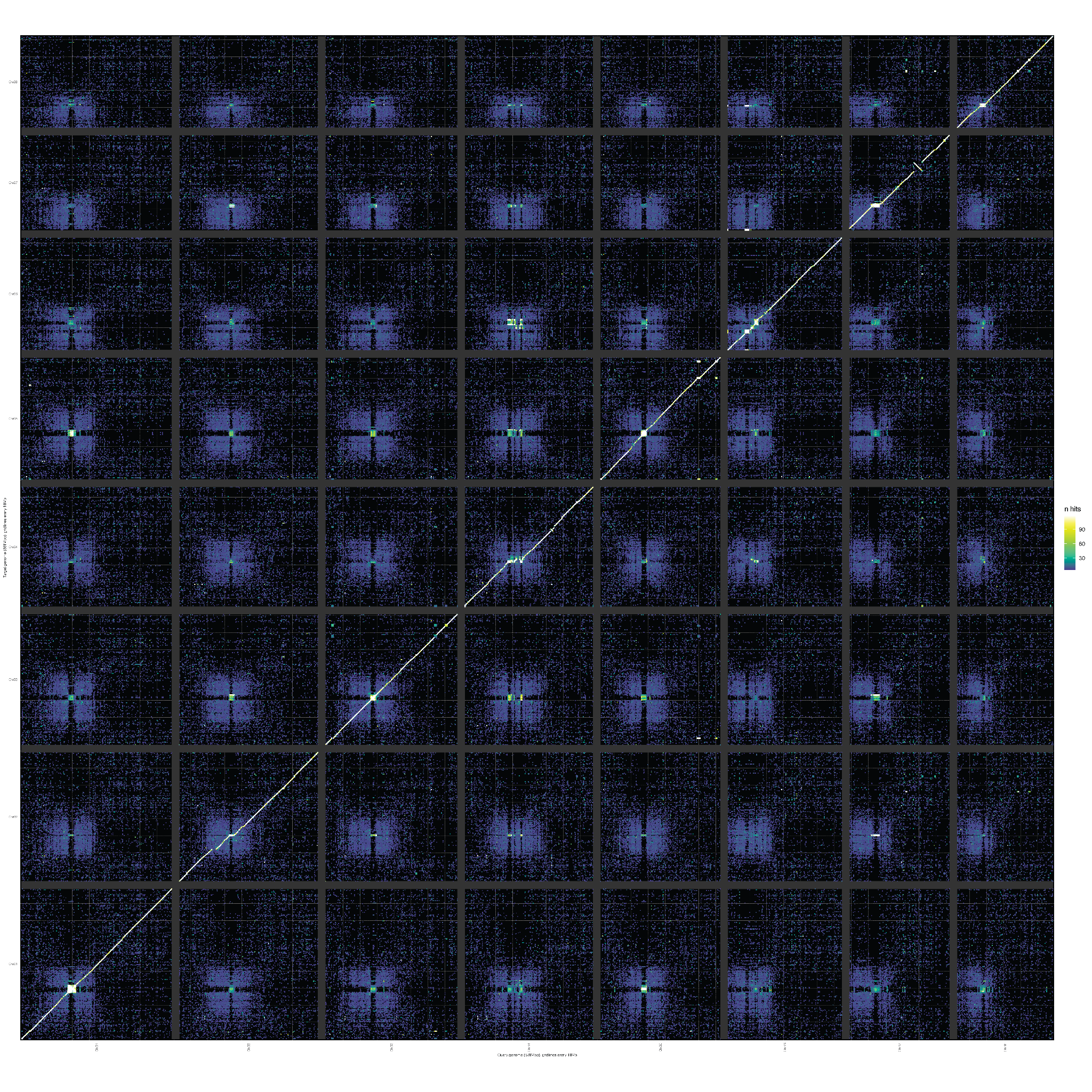

**Supplementary Figure 10.** Comparisons of haplotype assemblies for *C. fasciculata*. Regions of higher densities (lighter) show higher identities, indicating either conserved synteny or repetitive sequence.

**Supplementary Table 1.** List of genomes used in the initial gene family. For genera with multple species, these were incorporated into genus-level pangenes.

| **Genus** | **Species** | **Subfamily** | **Citation** | **doi** |
| --- | --- | --- | --- | --- |
| ***Acacia*** | *crassocarpa* | Caesalpinioideae | Massaro et al., 2023 | 10.1093/g3journal/jkad284 |
| ***Aeschynomene*** | *evenia* | Papilionoideae | Quilbé et al., 2021 | 10.1038/s41467-021-21094-7 |
| ***Cercis*** | *canadensis* | Cercidoideae | **This manuscript** |  |
| ***Chamaecrista*** | *fasciculata* | Caesalpinioideae | **This manuscript** |  |
| ***Dalbergia*** | *odorata* | Papilionoideae | Hong et al., 2020 | 10.1093/gigascience/giaa084 |
| ***Lablab*** | *purpureus* | Papilionoideae | Njaci et al., 2023 | 10.1038/s41467-023-37489-7 |
| ***Lens*** | *culinaris* | Papilionoideae | Ramsay et al., 2021 | 10.1101/2021.07.23.453237 |
| ***Lotus*** | *japonicus* | Papilionoideae | Sato et al., 2008 | 10.1093/dnares/dsn008 |
| ***Lupinus*** | *albus* | Papilionoideae | Hufnagel et al., 2020 | 10.1038/s41467-019-14197-9 |
| ***Phanera*** | *championii* | Cercidoideae | Lu et al., 2024 | 10.1111/tpj.16620 |
| ***Pisum*** | *sativum* | Papilionoideae | Kreplak et al., 2019 | 10.1038/s41588-019-0480-1 |
| ***Senna*** | *tomentosa* | Papilionoideae | Kang et al., 2020 | 10.1038/s41467-020-19681-1 |
| ***Sindora*** | *glauca* | Detarioideae | Yu et al., 2022 | 10.3389/fpls.2021.794830 |
| ***Trifolium*** | *pratense* | Papilionoideae | De Vega et al., 2015 | 10.1038/srep17394 |
| ***Vicia*** | *faba* | Papilionoideae | Jayakodi et al., 2023 | 10.1038/s41586-023-05791-5 |
| ***Arachis*** | *hypogaea* | Papilionoideae | Bertioli et al., 2019 | 10.1038/s41588-019-0405-z |
| ***Arachis*** | *stenosperma* | Papilionoideae | Bertioli et al., 2019 | 10.1038/s41588-019-0405-z |
| ***Arachis*** | *duranensis* | Papilionoideae | Bertioli et al., 2016 | 10.1038/ng.3517 |
| ***Arachis*** | *ipaensis* | Papilionoideae | Bertioli et al., 2016 | 10.1038/ng.3517 |
| ***Cicer*** | *arietinum* | Papilionoideae | Garg et al., 2021 | 10.1016/j.jare.2021.10.009 |
| ***Cicer*** | *echinospermum* | Papilionoideae | Cook et al., 2022 | GenBank GCA_002896215.2 |
| ***Cicer*** | *reticulatum* | Papilionoideae | Cook et al., 2022 | GenBank GCA_002896235.1 |
| ***Glycine*** | *max* | Papilionoideae | Schmutz et al., 2010 | 10.1038/nature08670 |
| ***Glycine*** | *soja* | Papilionoideae | Xie et al., 2019 | 10.1038/s41467-019-09142-9 |
| ***Glycine*** | *cyrtoloba* | Papilionoideae | Zhuang et al., 2022 | 10.1038/s41477-022-01102-4 |
| ***Glycine*** | *dolichocarpaD3* | Papilionoideae | Zhuang et al., 2022 | 10.1038/s41477-022-01102-4 |
| ***Glycine*** | *tomentella-D3* | Papilionoideae | Zhuang et al., 2022 | 10.1038/s41477-022-01102-4 |
| ***Glycine*** | *falcata* | Papilionoideae | Zhuang et al., 2022 | 10.1038/s41477-022-01102-4 |
| ***Glycine*** | *stenophita* | Papilionoideae | Zhuang et al., 2022 | 10.1038/s41477-022-01102-4 |
| ***Glycine*** | *syndetika* | Papilionoideae | Zhuang et al., 2022 | 10.1038/s41477-022-01102-4 |
| ***Medicago*** | *truncatula* | Papilionoideae | Tang et al., 2014 | 10.1186/1471-2164-15-312 |
| ***Medicago*** | *sativa* | Papilionoideae | Chen et al., 2020 | 10.1038/s41467-020-16338-x |
| ***Phaseolus*** | *acutifolius* | Papilionoideae | Moghaddam et al., 2021 | 10.1038/s41467-021-22858-x |
| ***Phaseolus*** | *lunatus* | Papilionoideae | Garcia et al., 2021 | 10.1038/s41467-021-20921-1 |
| ***Phaseolus*** | *vulgaris* | Papilionoideae | Schmutz et al., 2014 | 10.1038/ng.3008 |
| ***Vigna*** | *angularis* | Papilionoideae | Sakai et al., 2015 | 10.1038/srep16780 |
| ***Vigna*** | *radiata* | Papilionoideae | Ha et al., 2021 | 10.1002/tpg2.20121 |
| ***Quillaja*** | *saponaria* | Quillajaceae | Reed et al., 2023 | 10.1126/science.adf3727 |
| ***Arabidopsis*** | *thaliana* | Brassicaceae | Cheng et al., 2017 | 10.1111/tpj.13415 |
| ***Prunus*** | *persica* | Rosaceae | Verde et al., 2017 | 10.1186/s12864-017-3606-9 |
| ***Vitis*** | *vinifera* | Vitaceae | The French–Italian Public Consortium for Grapevine Genome Characterization, 2007 | 10.1038/nature06148 |

**Supplementary Table 2.** Classification of repetitive elements of *C. canadensis* and *C. fasciculata*

|  |  | ***C. canadensis*** | | | ***C. fasciculata*** | | |
| --- | --- | --- | --- | --- | --- | --- | --- |
| **Type** |  | **Number of elements** | **Length occupied (bp)** | **Percentage of sequence (%)** | **Number of elements** | **Length occupied (bp)** | **Percentage of sequence (%)** |
| **Retroelements** |  | 46,144 | 44,016,712 | 12.94% | 98,544 | 167,631,216 | 28.88% |
| **SINEs** |  | 330 | 66,257 | 0.02% | 16,321 | 3,305,483 | 0.57% |
| **Penelope** |  | - | - | 0.00% | 3,345 | 536,935 | 0.09% |
| **LINEs** |  | 12,009 | 7,197,792 | 2.12% | 13,915 | 7,156,358 | 1.23% |
|  | CRE/SLACS | - | - | 0.00% | - | - | 0.00% |
|  | L2/CR1/Rex | - | - | 0.00% | 33 | 37,051 | 0.01% |
|  | R1/LOA/Jockey | 1,889 | 3,253,041 | 0.96% | - | - | 0.00% |
|  | R2/R4/NeSL | - | - | 0.00% | - | - | 0.00% |
|  | RTE/Bov-B | 1,716 | 269,684 | 0.08% | 2,258 | 464,649 | 0.08% |
|  | L1/CIN4 | 8,332 | 3,627,676 | 1.07% | 11,624 | 6,654,658 | 1.15% |
| **LTR elements** |  | 33,805 | 36,752,663 | 10.80% | 68,308 | 157,169,375 | 27.08% |
|  | BEL/Pao | 447 | 51,271 | 0.02% | - | - | 0.00% |
|  | Ty1/Copia | 14,850 | 11,910,975 | 3.50% | 51,586 | 133,215,406 | 22.95% |
|  | Gypsy/DIRS1 | 17,270 | 22,898,626 | 6.73% | 14,764 | 23,075,603 | 3.98% |
|  | Retroviral | 173 | 71,013 | 0.02% | 917 | 210,672 | 0.04% |
| **DNA transposons** |  | 14,340 | 7,213,145 | 2.12% | 30,546 | 8,727,862 | 1.50% |
|  | hobo-Activator | 5,152 | 1,702,409 | 0.50% | 4,617 | 1,214,584 | 0.21% |
|  | Tc1-IS630-Pogo | 124 | 92,957 | 0.03% | 10,632 | 2,148,419 | 0.37% |
|  | En-Spm | - | - | 0.00% | - | - | 0.00% |
|  | MULE-MuDR | 4,628 | 3,265,000 | 0.96% | 5,211 | 2,843,971 | 0.49% |
|  | PiggyBac | - | - | 0.00% | - | - | 0.00% |
|  | Tourist/Harbinger | 1,313 | 584,355 | 0.17% | 1,834 | 436,523 | 0.08% |
|  | Other (Mirage, P-element, Transib) | - | - | 0.00% | - | - | 0.00% |
| **Rolling-circles** |  | 7,846 | 3,923,495 | 1.15% | 5,684 | 2,468,239 | 0.43% |
| **Unclassified** |  | 299,555 | 112,210,144 | 32.98% | 616,342 | 174,026,050 | 29.98% |
| **Total interspersed repeats** |  |  | 163,440,001 | 48.03% |  | 350,922,063 | 60.46% |
| **Small RNA** |  | 624 | 3,258,780 | 0.96% | 16,795 | 5,040,471 | 0.87% |
| **Satellites** |  | - | - | 0.00% | - | - | 0.00% |
| **Simple repeats** |  | 79,978 | 2,981,502 | 0.88% | 170,520 | 7,089,649 | 1.22% |
| **Low complexity** |  | 16,168 | 764,505 | 0.22% | 25,007 | 1,234,954 | 0.21% |

**Supplementary Table 3**. Whole-genome duplication and speciation peaks analysis by Ks value

| **Species 1** | **Species 2** | **WGD peak** | **speciation peak** |
| --- | --- | --- | --- |
| ***Bauhinia*** | ***Bauhinia*** | 0.25 |  |
|  | *Cercis* |  | 0.20 |
|  | *Medicago* |  | 0.90 |
|  | *Phaseolus* |  | 0.80 |
|  | *Quillaja* |  | 0.75 |
|  | *Senna* |  | 0.60 |
|  | *Sindora* |  | 0.65 |
| ***Cercis*** | ***Cercis*** | 2.00 |  |
|  | *Medicago* |  | 0.80 |
|  | *Phaseolus* |  | 0.70 |
|  | *Quillaja* |  | 0.65 |
|  | *Senna* |  | 0.50 |
|  | *Sindora* |  | 0.55 |
| ***Medicago*** | ***Medicago*** | 1.00 |  |
|  | *Phaseolus* | 0.90 | 0.75 |
|  | *Quillaja* |  | 1.10 |
|  | *Senna* |  | 0.90 |
|  | *Sindora* |  | 0.95 |
| ***Phaseolus*** | ***Phaseolus*** | 0.80 |  |
|  | *Quillaja* |  | 1.00 |
|  | *Senna* |  | 0.80 |
|  | *Sindora* |  | 0.90 |
| ***Quillaja*** | ***Quillaja*** | 0.30 |  |
|  | *Senna* |  | 0.80 |
|  | *Sindora* |  | 0.85 |
| ***Senna*** | ***Senna*** | 0.65 |  |
|  | *Sindora* |  | 0.70 |
| ***Sindora*** | ***Sindora*** | 0.60 |  |

**Supplementary Data 1.** Consensus sequences of candidate centromeric satellites for *C. fasciculata* and *C. Canadensis*.

>Cent-Cerca_104

AAAAAAATcTgAGTcAGAAAAAGGGAaGAAACGAAACAGGGTCCCGGgCCAaATATAGTGACTcACCAGGAAACtnTCCCnTCCCanATTcAGtgatntacaag

>Cent-Chafa_351

AAAACGAAGCAACAnGCATCTTCCCCTCAACTCTAACCTAAGATACCATTTAATTACTTGATTGTTGGACAAACCATGGGCGTGAGTGGCTTGGAACAACTTGAAACGGATCTTGGATGATAGGAAACGTTGTATGnTTGAAnCTTGnAnCAAGATTnGCCTCCATnnTATnACCTAnGGCCCnCCTnCnnCCCTCAACCTTGGTAATCACAACCTCCCCCAAGCTnAATCCTCTTCTAGGGAGGATGTCTAGGCTCTCAAGGTTTGAGATTGTGCTTCCACAAGCTCGAACTCCCTTTGATTGGCCATGGATATAGGGTAGACGATGTGGAGATGAGAAGAAGAGGAGAG

>Cent-Chafa_352

AAAACGAAGCAACATGCATCTTCCCCTCAACTCTAACCTAAGATACCATTTAATTACTTGATTGTTGGACAAACCATGGGCGTGAGTGGCTTGGAACAACTTGAAACCGATCTTGGATGATAGGAAACGTTGTATGCTTGAAGCTTGCAACAAGATTCGCCTCCATTGTATCAACTATGGACCACCCTCTACACTCTATCGTTGTGAATCACAACCTCCCCCAAGCTGAATCCTCTTCTAGGGAGGATGTCTAGGCTCTTAAGGTTTGAGATTGTGCTTCCACAAGCTCGAACTTCCTTTGATTGGCCATGGATATAGGGTAGACGATGTGGAGATGAGAAGATGAGGAGAG

>Cent-Chafa_355

AAACAAAGTTCGAGCTTGTGGGAGCACAATTTCAAACCTTAAGAGCCTAGACATCCTCCTCCCTnGAAGAGGATTCAGCTTGGGGGAGGTTGTTATTCACAACGTTAGAGTGTAGAGGGTGGTCCATAGTTGATACAATGGAGGCGAATCTTGTTGCAAGCTTCAAGCATACAACnTTTCCTATCATCCAAGATCGGTTTCAAGTTGTTCCAAGCCACTCACGCCCATGGTTTGTCCAACAATCAAGTAATTAAATGGTATnTTAGGTTAGAGTTGAGGGGAAGATGCATGTTGCTTCGTTTTCTCTCCTTATCTTCTCATCTCCACATCTTCTACCCTATATCCnTCGCCAATC
